# The Androgen Receptor and MYC synergise to modulate the synthesis of Siglec-7 ligands in prostate cancer

**DOI:** 10.1101/2025.10.15.682547

**Authors:** Adam Duxfield, Rebecca Garnham, Esme Hutton, Adam Dowle, Jamie Wills, Sara Luzzi, Ryan Nelson, Emma Lishman-Walker, Rianna Magee, Adriana Buskin, Anastasia C. Hepburn, Emirhan Tekoglu, Kaidan Maloney-Friar, Fiona Frame, Norman Maitland, Ann Hedley, Holly Henderson, Benjamin McCullough, Bharat Gowardhan, Kanagasabai Sahadevan, Stuart McCracken, Luke Gaughan, Rakesh Heer, Craig N. Robson, Kelly Coffey, Nathan Lack, Nathalie Signoret, Martin Fascione, Emma Scott

**Author notes:** These authors contributed equally (Adam Duxfield, Rebecca Garnham).

## Abstract

Glyco-immune checkpoints have recently been shown to be critical mediators of immunotherapy resistance across multiple cancer types. In clinical trials, immunotherapeutic treatments for prostate cancer have failed to elicit durable clinical responses. PCa progression is driven by transcriptional networks regulated by key transcription factors including the androgen receptor (AR) and the oncogene MYC. How this crossover between hormone and oncogene-driven signalling pathways regulates tumour glyco-immune checkpoints remains unclear. Here, we show that *O*-glycans are the major substrates for sialylation in prostate cancer and that sialyltransferases that have preferences for *O*-glycans are differentially regulated by androgens. We show that supraphysiological levels of androgens produce distinct glycopeptide profiles in prostate cancer cells compared with cells exposed to physiological androgens. Additionally, we identify a direct and coordinated role for AR and MYC in regulating ST3Gal1 and the synthesis of Siglec-7 ligands in prostate cancer. Both transcription factors converge to repress *ST3GAL1*, thereby limiting the generation of Siglec-7 ligands. These findings highlight a context-dependent, cooperative relationship between the AR and MYC in shaping the tumour sialome, linking hormonal signalling and oncogenic transcription to Siglec biology. Our study highlights how cell-type specific differences in transcriptional networks has important downstream effects for immune modulating glycans and has tumour specific clinical implications.

## Introduction

Prostate cancer (PCa) was responsible for deaths of over 100,000 men in Europe in 2022^1^. Its growth is fuelled by androgen receptor (AR) signalling, with the AR acting as a key hormone-activated transcription factor that controls cell survival, proliferation, and differentiation in the prostate epithelium^2,3^. While anti-androgen therapies initially provide clinical benefit, most patients ultimately develop resistance and progress to castration-resistant prostate cancer (CRPC), a stage of disease for which curative options remain elusive^3–5^.

Transcriptional reprogramming is a hallmark of PCa progression, with androgen signalling being the most notable and well-studied^6–8^. Alongside the AR, the *c-MYC* oncogene has emerged as a critical driver of PCa progression. MYC is frequently amplified in advanced prostate tumours and is known to reshape transcriptional, metabolic, and immune landscapes to support malignant growth^9–13^. Interestingly, while the AR and MYC can independently promote tumourigenesis, they also display transcriptional antagonism^14–16^. Some mechanisms have been identified, such as AR-mediated repression of MYC expression through super-enhancer remodelling, reciprocally, MYC disruption of AR target gene expression by modulating RNA polymerase II dynamics^15,17,18^. A middle ground has also been described, in which shared AR and MYC target genes are both positively regulated by each transcription factor^19,20^. The interplay between these two factors is complex and yet to be fully understood, but their balance, or imbalance, appears central to disease progression.

Recent evidence highlights glycosylation, particularly sialylation, as a key post-translational modification that can influence both cancer cell behaviour and immune evasion^21–24^. Sialyltransferases (STs) are enzymes that attach terminal sialic acids to glycoproteins and glycolipids, generating sialoglycans that can engage immunosuppressive Siglec receptors on myeloid cells. This interaction promotes tumour evasion and ultimately blunts anti-tumour immunity. In PCa, sialylation patterns are dynamic and have been shown to be modulated by androgens ^25–33^.

Among the twenty STs, three enzymes have been broadly studied in PCa. ST3Gal1, which generates α2,3-linked sialoglycans, is upregulated in prostate tumours with low AR activity and has been shown to drive Siglec-7 and Siglec-9 ligand formation, promoting immune evasion^30,34^. In contrast, ST6Gal1 and ST6GalNAc1, responsible for α2,6-linked sialylation, are positively regulated by the AR and support tumour growth and survival^28,29,35^. Collectively, ST3Gal1, ST6Gal1 and ST6GalNAc1 shape the PCa sialome.

While AR regulation of STs has been broadly described, nuances in AR-ST biology and the contribution of MYC remains poorly understood. Recent studies in other cancer types, including T-cell acute lymphoblastic leukaemia suggest that MYC can transcriptionally upregulate STs such as ST6GalNAc4, thereby enhancing sialoglycan-mediated immune suppression^36^. Whether MYC exerts similar control in PCa, especially in the context of AR antagonism, is currently unclear.

In this study, we demonstrate that physiological and supraphysiological levels of androgens produce distinct sialoglycan patterns in PCa and that both the AR and MYC independently regulate key STs reshaping the prostate cancer sialome. We show that the two transcription factors cooperate to repress ST3Gal1 and that the AR-MYC-ST3Gal1 signalling cascade alters Siglec-7 sialoglycan ligands. We describe a complex interplay between the AR and MYC which reveals a previously unrecognised axis linking hormonal signalling, oncogenic transcription, and tumour glycosylation. Our findings reveal a coordinated, context-dependent regulation of the sialome with implications for glyco-immune checkpoint targeting in advanced disease.

## Methods

### Cell culture

LNCaP cells were cultured in RPMI-1640 with 10% fetal calf serum (FCS) and 1% penicillin– streptomycin. VCaP cells were cultured in DMEM with 10% FCS and 1% penicillin– streptomycin. TRAMP-C2 cells were maintained in DMEM supplemented 0.005 mg/mL bovine insulin, 10 nM dehydroisoandrosterone, 5% FCS, and 5% Nu-Serum IV. All cells underwent regular mycoplasma testing.

### Bioinformatic analysis of publicly available datasets

Publicly available transcriptomic datasets were accessed using cBioPortal^37^ or camcAPP (https://bioinformatics.cruk.cam.ac.uk/apps/camcAPP/). Previously published datasets were downloaded from GEO. MYC activity scores were calculated as described by *Schaafsma et al* ^38^. Ecotyper analysis was performed on data available in the TCGA-PRAD dataset accessed using cBioPortal^37^. Data was uploaded and analysed using Ecotyper (https://ecotyper.stanford.edu/)^39^.

Raw data for AR and MYC ChIP-seq was downloaded from the GEO dataset GSE73994^15^ using sra-tools v.3.0.10. Quality of the reads was checked using fastqc v.0.12.1 before aligning them to the human genome GRCh38 (Ensembl 114) using bowtie v.1.3.1 with default parameters. SAM files were converted to BAM using samtools v.1.14 and peaks were detected using MACS2 v.2.2.7.1 with a q-value cutoff of 0.01. For visualisation on the IGV genome browser, replicate BAM files were merged with samtools and converted to bigWig using bedtools v.2.30.0 and bedGraphToBigWig v.2.10. Similarly, peak signals shown were obtained by merging MACS2 narrowPeak files using bedtools.

scRNA-seq cluster annotations and gene expression values for primary prostate cancer cells were obtained from the authors of Song et al, 2022 as a Seurat R object. Gene and cluster UMAPs were plotted using the Seurat FeaturePlot and DimPlot functions, respectively.

### Flow cytometry for Siglec ligands

1x10^6^ cells were collected and washed in cold PBS. Siglec-Fc chimeras were purchased from R&D Systems and used at a concentration of 4 µg/mL. Siglec-Fc were pre-complexed with an anti-human-PE secondary antibody for 30 minutes on ice prior to incubation with cells for 30 minutes. Antibodies used and their concentrations can be found in supplemental Table 1.

Cells were then washed three times in PBS and analysed using a FACSymphony A5 cytometer.

### Tissue co-immunofluorescence

Formalin-fixed, paraffin-embedded (FFPE) human PCa tissue sections were dewaxed, rehydrated through a graded alcohol series, and washed with TBS. Antigen retrieval was performed by heating slides in Tris-EDTA buffer at 121 °C for 15 minutes. Slides were then blocked with 10% goat serum for 1 hour at room temperature and incubated overnight at 4 °C with primary antibodies diluted in blocking solution. After washing with TBS-T, slides were incubated with either goat anti-rabbit Alexa Fluor 647 or donkey anti-mouse Alexa Fluor 488 secondary antibodies for 60 minutes at room temperature. Slides were washed in TBS-T and mounted using ProLong Gold Antifade mountant. Antibodies used and their concentrations can be found in supplemental Table 1. Imaging was performed using a ZEISS AxioImager 2 microscope using ZEN 2.6 software.

### Glycopeptide analysis

For samples preparation, pellets from 10 x 10^6^ LNCaP cells were resuspended in 1 ml of 5% SDS lysis buffer (5% SDS, 100 mM Tris pH 8.0, 50 mM DTT in double distilled H_2_O). Cells were passed multiple times through a 21-gauge needle, then heated at 95°C for 5 min. Lysates were clarified by centrifuging for 10 min at 10,000 rpm and prepared for glycoproteomic analysis using an S-trap procedure^40^.

Sample were lysed with aqueous 5% (v:v) sodium dodecyl sulfate, 100 mM triethylammonium bicarbonate (TEAB) and a 10 µg aliquot of protein lysate taken for analysis. Protein was reduced with 5.7 mM tris(2-carboxyethyl)phosphine and heating to 55°C for 15 mins before alkylation with 22.7 mM methyl methanethiosulfonate at room temperature for 10 mins. Protein was acidified with 6.5 μL of aqueous 27.5 % (v:v) phosphoric acid then precipitated with dilution seven-fold into 100 mM TEAB 90% (v:v) methanol. Precipitated protein was captured on S-trap (Profiti – C0-micro) and washed five times with 165 µL 100 mM TEAB 90% (v:v) methanol before digesting with the addition of 20 µL 0.1 µg/µL Promega Trypsin/Lys-C mix (V5071) in aqueous 50 mM TEAB and incubation at 47 °C for 2h. Peptides were recovered from S-trap by spinning at 4000 g for 60 s. S-traps were washed with 40 µL aqueous 0.2% (v:v) formic acid and 40 µL 50% (v:v) acetonitrile:water and washes combined with the first peptide elution. Peptide solutions were dried in a vacuum concentrator then resuspended in aqueous 0.1% (v:v) formic acid.

A 1 µg aliquot of each peptide solution was loaded onto EvoTip Pure tips for desalting and as a disposable trap column for nanoUPLC using an EvoSep One system. A pre-set EvoSep 15 SPD gradient (from Evosep One HyStar Driver 2.3.57.0) was used with a 15 cm EvoSep C_18_ Performance column (15 cm x 150 μm x 1.5 μm).

The nanoUPLC system was interfaced to a timsTOF HT mass spectrometer (Bruker) with a CaptiveSpray ionisation source (Source). Positive PASEF-DDA, ESI-MS and MS^2^ spectra were acquired using Compass HyStar software (version 6.2, Bruker). Instrument source settings were: capillary voltage, 1,700 V; dry gas, 3 l/min; dry temperature; 180°C. Spectra were acquired between *m/z* 100-1,700. TIMS settings were: 1/K0 0.6-1.60 V.s/cm^2^; Ramp time, 100 ms; Ramp rate 9.42 Hz. Data dependant acquisition was performed with 10 PASEF ramps and a total cycle time of 1.17 s. An intensity threshold of 2,500 and a target intensity of 20,000 were set with active exclusion applied for 0.4 min post precursor selection. Collision energy was interpolated between 20 eV at 0.6 V.s/cm^2^ to 59 eV at 1.6 V.s/cm^2^.

Data were searched using FragPipe (v23.0) against the human subset of UniProt appended with common proteomic contaminants ^41^. Search parameters specified trypsin protease with up to two missed cleavage positions allowed. Mass tolerance was set to 15 ppm for MS1 and MS2 matching. Beta-methylthiolation (45.9877Da) of was set as fixed modifications. Oxidation (15.995 Da) was selected as a variable modification. Additionally, all built-in O-glycan and N-glycan modifications within the pre-set glyco-O-HCD and glyco-N-HCD workflows of FragPipe were considers as variable modifications within MSFragger (v4.2). Validation was performed using ProteinProphet and PeptideProphet within Philosopher (v5.1.1). Searches were run at 1% FDR as assessed empirically against a reverse database search. Site localisation probabilities were calculated with PeptideProphet. Intensities from accepted glycopeptides were extracted using IonQuant (v1.11.9).

### Quantitative PCR

Cells were harvested and RNA was extracted using a RNeasy® Mini Kit (Qiagen, 74104) in accordance with the manufacturers protocol. cDNA synthesis was performed on 500 ng of RNA using the M-MLV Reverse Transcriptase (Thermo Fisher Scientific, 28025013). Quantitative PCR (qPCR) was performed in triplicate using SYBR® Green PCR Master Mix (Applied Biosystems™, 4309155) and analysed on the Quant Studio 7 Flex Real-Time PCR System (Thermo Fisher Scientific, 4485701). Gene expression values were normalised to the average of two housekeeping genes: Actin and β-tubulin. All primer sequences can be found in supplemental Table 2.

### Western blotting

Protein was extracted using 2x SDS lysis buffer containing DENERASE® (Avantor, 20804-100K) at a working concentration of 0.2 kU/mL for 10 minutes on ice. The protein lysates were denatured by heating at 95°C for 5 minutes, prior to being separated on a 10% acrylamide gel (Bio-Rad, 4561033) at 180 V. The gels were then transferred onto a nitrocellulose membrane using the Bio-Rad Trans-Blot® Turbo System according to the manufacturer’s instructions, using the High MW protocol. The membranes were then incubated in 5 % skimmed milk in TBS-Tween for 1 hour with agitation. Followed by an incubation with primary antibody overnight at 4 °C with rotation. Membranes were washed 3 times for 5 minutes in TBS-Tween (RT) and the appropriate secondary antibodies which were either conjugated to horseradish peroxidase (HRP) or fluorophore were added to the membrane for 1 hour at RT with agitation. After the incubation period the membranes were washed twice with TBS-Tween for 5 minutes at RT, followed by a final wash in TBS for 5 minutes at RT. For membranes incubated with HRP conjugated secondaries the membrane was incubated in enhanced chemiluminescence (ECL solution for 2 minutes) and then visualised using a Li-Cor Odyssey® Fc 2800 Western Blot Imaging System. Membranes incubated with fluorophore conjugated secondaries were imaged directly without any additional steps to enhance visualisation. The approximate size of the target protein was assessed by comparison against the PageRuler™ Plus pre-stained protein ladder (Thermo Fisher Scientific, 26619). Each western blot shows technical replicates from 1 biological experiment. All western blot experiments have been repeated a minimum of 3 times representing 3 independent biological replicates. All replicates and uncropped western blots can be found in supplemental figure 7. Information on all the antibodies used can be found in supplemental Table 1.

### Lectin flow cytometry

MAL-II expression was assessed using biotinylated MAL-II lectin (Vector Laboratories, B-1305-2), followed by detection with streptavidin-conjugated Alexa Fluor 647. A total of 1 × 10^6^ cells were pelleted by centrifugation, washed with cold PBS and resuspended in FACS buffer (PBS-5% FBS). MAL-II lectin was pre-complexed with AF647-streptavidin on ice for 30 minutes. Cells were incubated with pre-complexed MAL-II lectin (at 2 µg/mL) for 30 minutes at 4 °C. SNA expression was assessed using a FITC-conjugated SNA lectin (diluted 1:1000 in PBS). 1 × 10^6^ cells were incubated with the FITC-SNA lectin for 30 minutes on ice and washed three times. Samples were analysed on a FACSymphony A5 flow cytometer.

### Animal models

All animal experiments were conducted in accordance with the UK Animals (Scientific Procedures) Act 1986 and approved under a UK Home Office licence (PC02CF4AB). Procedures adhered to the ARRIVE guidelines, and all relevant ethical regulations for animal use were followed. Ethical approval was granted by the Newcastle University Animal Welfare and Ethical Review Board (AWERB). Mice were housed under standard conditions with ad libitum access to food and water and maintained on a 12-hour light–dark cycle.

Male C57BL/6 mice (7 weeks old) were obtained from Charles River. 8-week-old mice were injected in the right flank with 2 × 10⁶ TRAMP-C2 cells. Mice were allowed to establish tumours prior to random allocation into treatment groups. Mice received 20 mg/kg enzalutamide or DMSO vehicle by oral gavage once daily at the indicated time points.

### Lectin and Siglec-ligand immunofluorescent staining of tissue

FFPE sections were dewaxed for 10 minutes in histoclear and rehydrated in graded alcohols. Antigen retrieval was performed in 1 mM EDTA at 90°C for 20 minutes. Non-specific binding was blocked using 10% goat serum in PBS-T for 1 hour at RT, washed and further blocked for 1 hour in 10% casein. Sections were washed 3 times in PBS-T and MAL-II or SNA lectin (diluted 1:1000) or Siglec-FC-chimeras were added and sections incubated overnight at 4°C. For MAL-II staining slides were washed 3 times in PBS-T and then incubated with a AF647-streptavidin for one hour at room temperature in the dark. For Siglec-ligand chimeras a AF647-anti-human-Fc was used as a secondary antibody. Sections were again washed and incubated in Hoechst for 15 minutes. Fluorescent images were obtained with a ZEISS Axio Imager 2 (Carl Zeiss, Germany) using the ZEN 2.6 software.

The level of cellular fluorescence from immunofluorescent images was determined using ImageJ2 v2.9.0. Fluorescent intensity was calculated using the Freehand ROI tool to measure the fluorescent intensity at the cellular level using 5 representative images per condition. For each representative image, five different regions of interest of cell clusters (5 cells per cluster) were measured to calculate integrated density. Corrected total cell fluorescence was calculated by subtracting background fluorescence measurements at regions neighbouring measured cells. Corrected total cell fluorescence (CTCF) = Integrated Density – (area of cell cluster x mean fluorescence of background readings).

### Ex vivo precision cut tissue slicing

Following robotic radical prostatectomy precision cut tissue slices were grown as described previously ^42^ with a slight modification, tissue was cultured in transwells instead of on sponges. Tissue slices were randomly assigned to wells and allowed to recover overnight. The following day treatments were applied to a minimum of 3 slices per condition, and thereafter culture medium was replenished every 24 hours. Darolutamide was added to a final concentration of 10 µM or the equivalent concentration of DMSO (vehicle control). After 48 hrs of treatment tissue slices were washed in PBS and either snap-frozen or fixed in 4% paraformaldehyde at room temperature. After fixation tissue was transferred to 70% ethanol prior to processing with the (Thermo Fisher Scientific Excelsior tissue processor) and embedding in paraffin wax.

### Stimulation and drug treatment of prostate cancer cells

For androgen stimulation experiments the LNCaP cells were cultured in steroid-depleted media (media supplemented with dextran charcoal-stripped foetal bovine serum) for 72 hours. Following this either 1 nM or 10 nM R1881 or 10 nM DHT was added for 24 hours. The cells were then harvested for protein, RNA and for flow cytometry analysis.

Enzalutamide was used at a final working concentration of 10 µM for all experiments. Darolutamide (ODM-201) was used at a final working concentration of 10 µM for all experiments. The cells were treated for a period of 48 hours before being harvested for protein, RNA and for flow cytometry. Control cells were treated with an equivalent concentration of DMSO as a vehicle control.

### Sh/siRNA knockdown studies

The generation of cell lines with stable knockdown of *ST3GAL* were performed by lentiviral transduction using a multiplicity of infection (MOI) of 5 for both shControl and shST3GAL1 as described previously^30^. Successfully transduced cells were selected using 2 µg/mL puromycin.

A transient knockdown of c-MYC was generated by transfecting cell lines with, scrambled non-silencing (control), MYC #1 (sequence: CGUCCAAGCAGAGGAGCAA[dT][dT]) and MYC #2 (sequence: GCUUGUACCUGCAGGAUCU[dT][dT])^42^ siRNA using the RNAiMAX transfection reagent according to the manufacturers protocol. Cells were treated at 60-70% confluency and incubated for a period of 48 hours before being harvested.

### Human tissue sample ethics

FFPE tissue samples were collected with permission from Castle Hill Hospital (Cottingham, Hull) (Ethics Number: 07/H1304/121) and tissue for *ex vivo* tissue slicing from Sunderland Hospital (REC reference: 23/WM/0022). Approval was given by the Local Research Ethics Committees. Patients gave informed consent, and all patient samples were anonymised. All ethical regulations relevant to human research participants were followed.

### Data availability

All mass spectrometry data and glycopeptide database search results are deposited in MassIVE (MSV000099433) and referenced in ProteomeXchange (PXD069255).

Data are currently password protected but can be accessed via: ftp://MSV000099433@massive-ftp.ucsd.edu

### Statistics and reproducibility

Statistical analyses were performed using GraphPad Prism 8 (GraphPad Software, Inc., San Diego, CA, USA). Statistical significance is shown as * *p* < 0.05, ** *p* < 0.01, *** *p* < 0.001 and **** *p* < 0.0001. Individual sample sizes are defined in individual figure legends.

## Results

### Sialyltransferase expression in prostate cancer

Tumour associated changes in sialoglycan patterns are governed by several mechanisms, including differential expression of sialyltransferases (STs) and changes to enzymes which are critical for the biosynthesis of sialic acid^43^. Sialic acid biosynthesis is a multistep process whereby Uridine diphosphate *N*-acetylglucosamine (UDP-GlcNAc) is converted to *N*-Acetylneuraminic acid (Neu5Ac), better known as sialic acid, and is catalysed by several enzymes, including UDP-*N*-acetylglucosamine 2-epimerase (GNE), *N*-acetylneuraminate synthase (NANS) and *N*-acylneuraminate-9-phosphatase (NANP) (**figure 1a**). Gene expression of two of these critical enzymes, *GNE* and *NANS*, is significantly increased in prostate cancer patients (N=492), when compared with normal prostate tissue (N = 152) (**figure 1b**). Next, we profiled transcript levels of all 20 STs in both The Cancer Genome Atlas (TCGA) Prostate Adenocarcinoma (PRAD) cohort (N= 489) and an independent cohort of metastatic castrate resistant PCa patients (N = 208)^44^. Across all datasets, ST6Gal1, ST6GalNAc1, ST3Gal1 and ST3Gal4 are the most abundantly expressed STs in prostate tumours (**figure 1c and supplemental figure 1a**).

**Figure 1.**
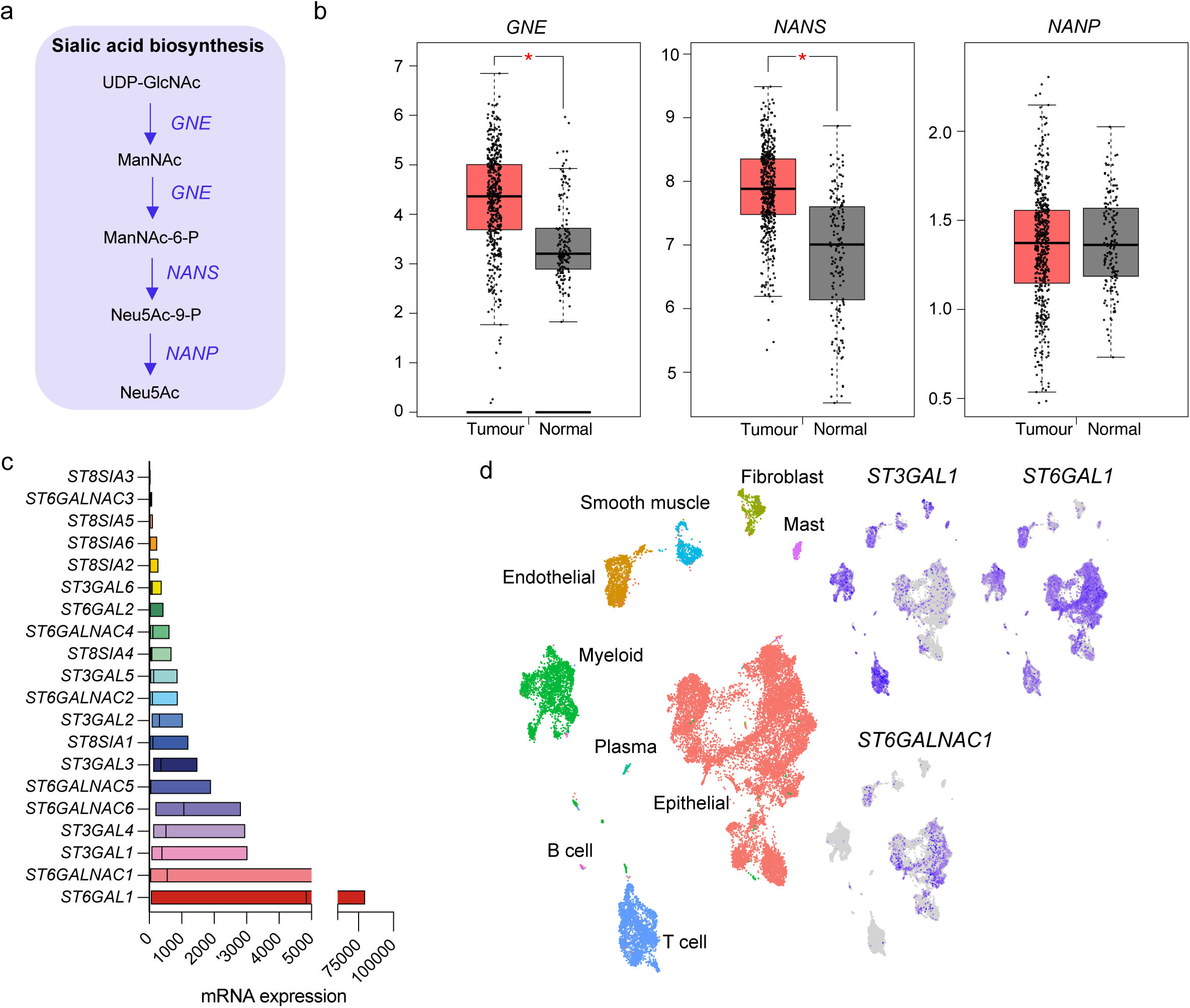
Sialyltransferase expression in prostate cancer. (**a**) A schematic of the sialic acid biosynthesis pathway. This process is initiated by the enzyme UDP-N-acetylglucosamine 2-epimerase/N-acteylmannosamine kinase (GNE) which converts UDP-GlcNAc, the main product of the hexosamine biosynthetic (glucose metabolism) pathway to form N-acetylmannosamine (ManNAc). GNE then phosphorylates ManNAc to produce N-acetylmannosamine-6-phosphate (ManNAc-6-AP). The second enzyme involved in this process is Neu5Ac synthase (NANS) which forms Neu5Ac-9-P in a condensation reaction of ManNAc-6-P with phosphoenolpyruvate (PEP). The final step of this pathway involves the removal of 9-phosphate by the enzyme N-acylneuraminate-9-phosphatase (NANP) to produce Neu5Ac (sialic acid). (**b**) mRNA expression of the three-core sialic acid biosynthesis enzymes (*GNE*, *NANS* and *NANP*) in The Cancer Genome Atlas (TCGA) Prostate Adenocarcinoma (PRAD) cohort (N=492) compared with normal prostatic tissue (N=152) Data accessed and plotted using GEPIA (https://gepia.cancer-pku.cn/) (**c**) mRNA levels of all twenty sialyltransferases in transcriptomic data from the TCGA PRAD cohort (Accessed through cBioPortal, N=489). (**d**) Uniform manifold approximation and projection (UMAP) showing clustering of cell types in single-cell transcriptomic analysis of human primary prostate tumours from a previously published dataset ^45^. Cell specific gene expression of *ST3GAL1*, *ST6GAL1* and *ST6GALNAC1* is shown.

Focusing on the most abundantly expressed STs, single-cell transcriptomics was employed to examine their expression across cell types within the PCa TME. Single-cell expression profiles from PCa tumours revealed that *ST3GAL1* and *ST6GAL1* are widely expressed throughout the prostate TME, including in the epithelium and stroma (**figure 1d**)^45^. Whilst expression of *ST6GALNAC1* is restricted to the epithelial compartment and endothelial cells. Our data show that *ST3GAL1*, *ST6GAL1* and *ST6GALNAC1* are the most abundantly expressed STs in prostate tumours and are broadly expressed throughout the prostate TME.

### Siglec receptors are expressed on tumour associated macrophages in prostate tumours

Sialic acid-binding immunoglobulin-like lectins (Siglec) receptors are broadly expressed immunoreceptors. Several contain immune tyrosine-based inhibitors motifs (ITIMs) which function to suppress the immune response. Sialoglycans synthesised by sialyltransferases serve as Siglec ligands. As such, the Siglec receptors are responsible for many of the pro-tumour functions of sialoglycans. Several of the Siglec receptors have been linked to ST3Gal1, ST6Gal1 and ST6GalNAc1. We previously reported that ST3Gal1 synthesises Siglec-7 and Siglec-9 ligands ^30^. ST6Gal1 has been shown to synthesise CD22, CD33, and Siglec-10 ligands and ST6GalNAc1 has been linked to the synthesis of Siglec-7 and Siglec-15 ligands. We probed Siglec ligand expression in LNCaP prostate cancer cells using Siglec-Fc reagents made of the extracellular portion of the Siglec receptor fused to a human IgG Fc domain (flow cytometry gating strategy shown in **supplemental figure 2a**). LNCaP PCa cells display relatively high levels of Siglec-7, Siglec-9 and Siglec-15 ligands, with low levels of CD22, CD33 and Siglec-10 ligands (**figure 2a and supplemental figure 2b**). We have previously confirmed that Siglec-7/9 ligands co-localise with α-methylacyl-CoA racemase (AMACR), indicating that they are expressed in glandular structures in prostate tumours. We now confirm that CD33, Siglec-10 and Siglec-15 ligands co-localise with AMACR and are expressed in cancerous prostate glands (**figure 2b**). Although low levels of ligands for CD22, which preferentially bind *N*-linked sialoglycans synthesised by ST6Gal1, can be found in AMACR^+^ glands, the majority are displayed in stromal regions of prostate tumours^46–48^. We observed similar stromal staining patterns for Siglec-15 ligands.

**Figure 2.**
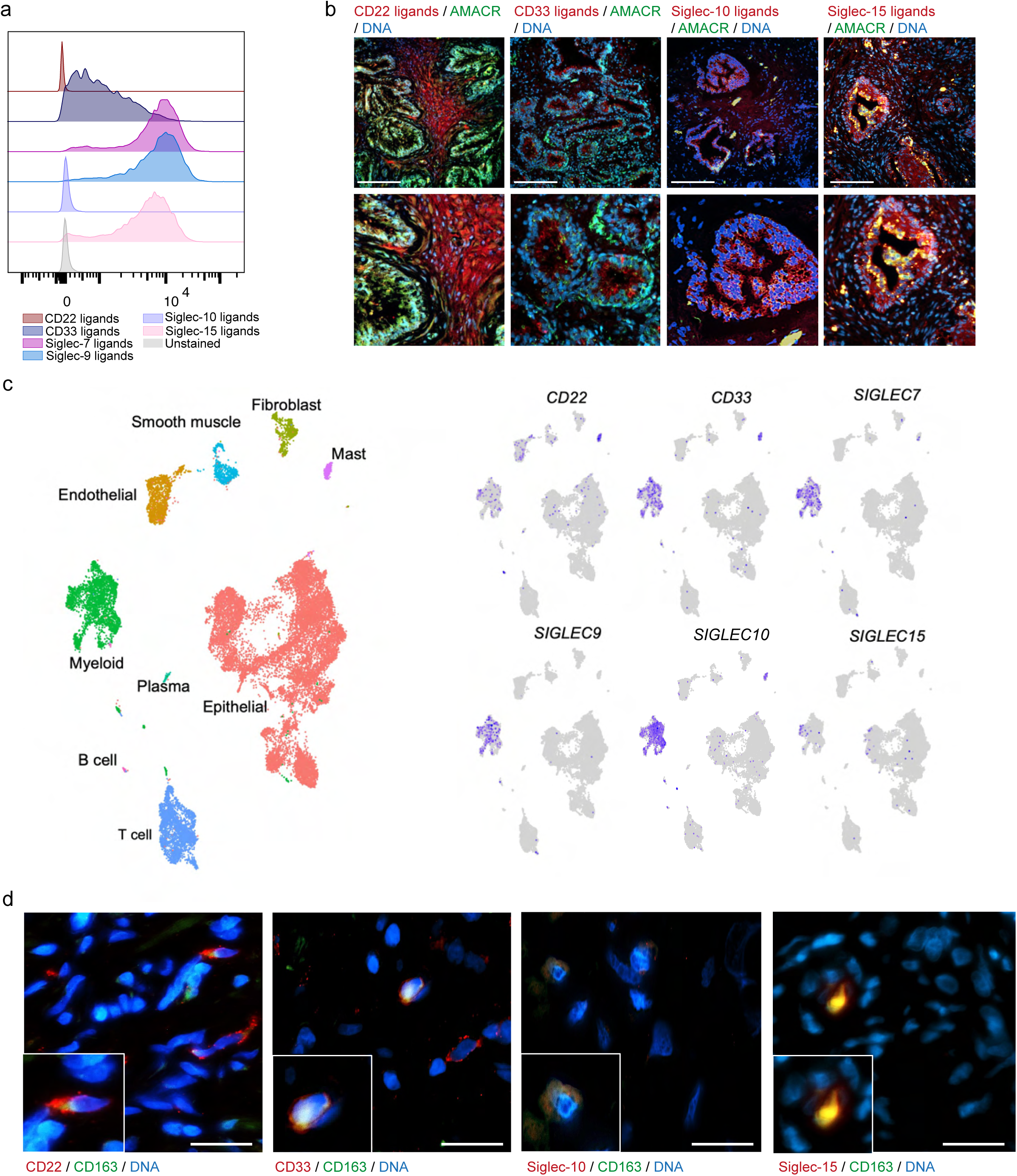
Siglec profiling in prostate cancer. (**a**) Quantification of the basal expression of CD22, CD33, Siglec 7, Siglec 9, Siglec 10 and Siglec 15 immunosuppressive ligands in LNCaP cells using Siglec-Fc chimeras. Representative histogram of *N* = 3 biologically independent samples. (**b**) The ligands for CD22, CD33, Siglec-10 and Siglec-15 (red) co-localised with alpha-methylacyl-CoA racemase (AMACR) (green) in prostate cancer patient biopsies using dual immunofluorescence. Images were prepared using a ZEISS Axio Imager2 microscope with a x20 and x40 objective. Scale bar = 150 µm (**c**) Uniform manifold approximation and projection (UMAP) showing clustering of cell types in single-cell transcriptomic analysis of human primary prostate from a previously published dataset ^45^. Gene expression of six Siglec immunoreceptors (*CD22*, *CD33*, *SIGLEC 7*, *SIGLEC 9* and *SIGLEC 15*) are shown. (**d**) The CD22, CD33, Siglec-10 and Siglec-15 immunoreceptors (red) co-localised with CD163 (green) in prostate cancer patient biopsies using dual immunofluorescence. Images were prepared using a ZEISS Axio Imager2 microscope with a x20 objective. Scale bar = 20 µm.

Siglec receptors are known to be expressed throughout the immune system, with many being predominantly expressed by myeloid cells. Analysis of previously published single cell-RNA sequencing of primary prostate tumours confirms that *CD33, SIGLEC7, SIGLEC9*, *SIGLEC10* and *SIGLEC15* are mainly expressed on myeloid populations, in line with literature on other cancers^45,49–58^ (**figure 2c**). CD22 has often been described as a B-cell restricted marker^59–61^. Our data confirms *CD22* expression on B cells but indicates that *CD22* is also expressed on some myeloid and endothelial cells in prostate tumours. Myeloid cells have recently been shown to be key determinants of immune suppression in prostate tumours and high numbers of CD163^+^ tumour associated macrophages (TAMs) in prostate tumours are predictive of poor clinical outcomes^62–64^. We have previously shown that both Siglec-7 and Siglec-9 are expressed on TAMs in prostate tumours. Using immunofluorescence co-staining, we confirm that in prostate cancer CD22, CD33, Siglec-10 and Siglec-15 are expressed on CD14^+^ myeloid cells (**supplemental figure 2c**) and CD163^+^ TAMs (**figure 2d**). These data confirm that the Siglec ligands that are made by ST3Gal1, ST6Gal1 and ST6GalNAc1 are expressed in prostate tumours, and that the Siglec receptors which recognise these sialoglycans are expressed by CD163^+^ TAMs which are poorly prognostic in prostate cancer patients.

### Sialic acid is predominantly found on O-glycans in prostate cancer cells

Sialoglycans have previously been shown to be modulated by androgens in PCa^28,29^. We performed glycopeptide analysis to produce an unbiased profile of glycopeptides in LNCaP cells stimulated with 1 nM R1881 to model physiological androgen levels. Across both conditions we identified 598 glycopeptides of which 7.3% were sialylated (**figure 3a**). When looking at all identified glycopeptides, 42.5% were *O*-glycans, whilst 90.0% of sialylated glycopeptides were *O*-glycans (**figure 3b**). When looking at the 44 sialylated glycopeptides, the most abundant glycan structures were common *O*-glycan structures; the Sialyl-T antigen and the di-sialyl-T antigen (**figure 3c**). When LNCaP cells were treated with synthetic androgens, the composition of sialylated glycans changed, with a loss of complex sialylated glycopeptides and an increase in glycopeptides containing the Sialyl-T antigen (**figure 3d**). ST3Gal1 primarily sialylates galactose in core-1-*O*-glycans forming the Sialyl-T structure. ST6GalNAc1 can sialylate the underlying GalNAc residue forming the disialyl-T structure or sialylate lone GalNAc residues forming the Sialyl-Tn *O*-glycan. ST6Gal1 is responsible for the sialylation of galactose on *N*-glycan structures. (**figure 3e**). These data show that O-glycans are the largest source of sialic acid in PCa cells.

**Figure 3.**
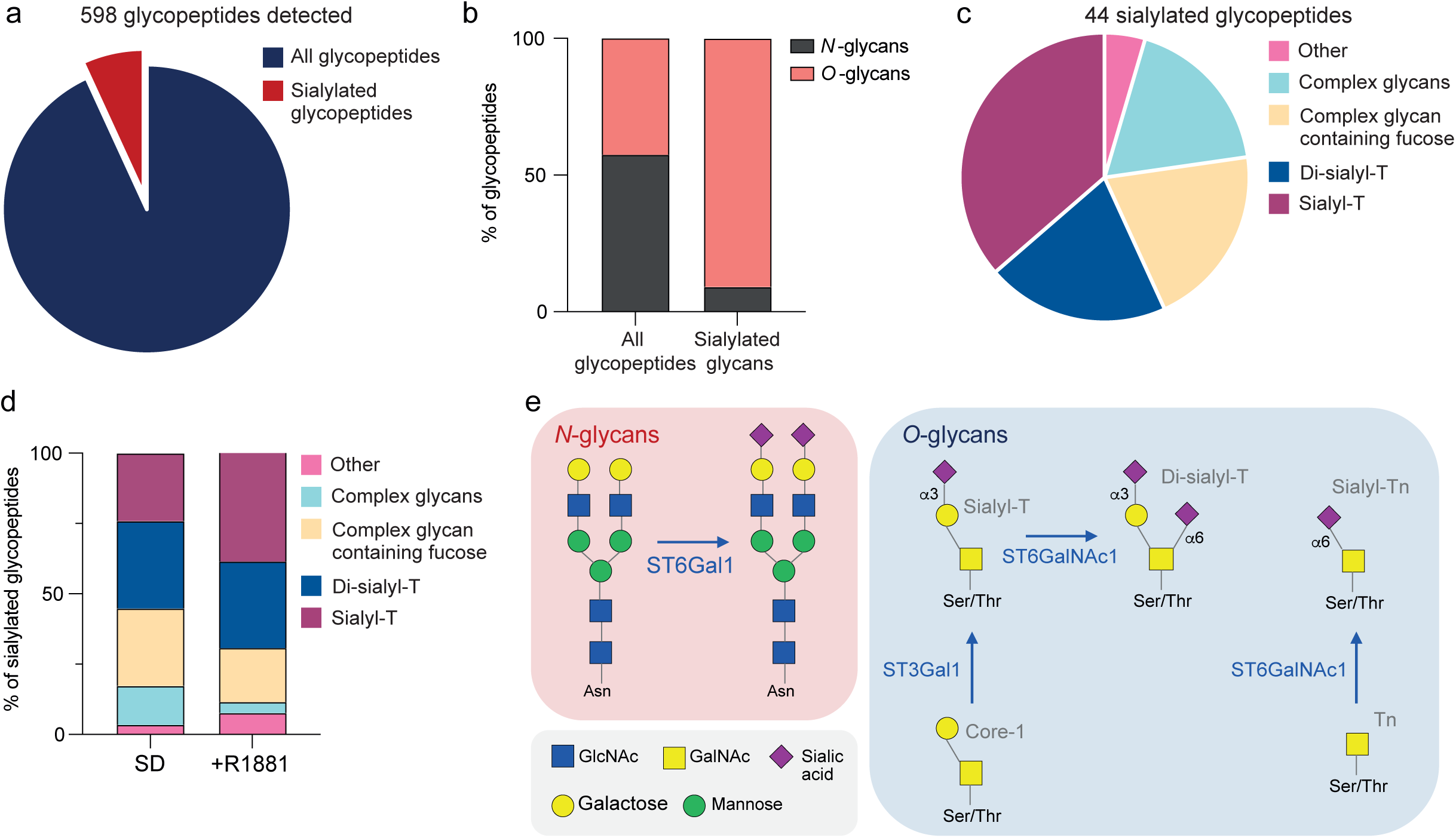
Sialic acid is predominantly found on *O*-glycans in prostate cancer cells. (**a**) Pie chart showing the percentage of glycopeptides that are sialylated from 598 glycopeptides detected in LNCaP cells. (**b**) Stacked bar chart showing the percentage of *N*- and *O*-glycans in all glycopeptides, and sialylated glycopeptides in LNCaP cells. (**c**) Pie chart showing the percentage of each glycan species detected in 44 sialylated glycopeptides in LNCaP cells. (**d**) Stacked bar chart showing the percentage of each sialoglycan species in LNCaP cells that were steroid depleted and LNCaP cells treated with 1 nM R1881 for 24 hours. (**e**) Schematic showing how ST3Gal1, ST6Gal1 and ST6GalNAc1 contribute to the synthesis of *N*- or *O*-linked sialoglycans.

### STs with preferences for O-glycans are modulated by androgens

Previously, both ST6Gal1 and ST6Gal1NAc1 have been reported to be positively regulated by androgens, whilst ST3Gal1 activity has been shown to be negatively correlated with androgen receptor activity. To confirm this, we treated LNCaP cells with 10 nM dihydrotestosterone (DHT) or 1 nM of the synthetic androgen R1881 to model physiological levels of androgens found within the prostate^65^. We analysed gene expression of the 20 STs and found that ST genes which preferentially sialylate *O*-glycans are significantly altered in response to AR stimulation (**figure 4a**). *ST6GALNAC1* and *ST6GALNAC3* are both significantly upregulated in response to both DHT and R1881 with the greatest increase observed for *ST6GALNAC1* (**figure 4a**). *ST3GAL1, ST3GAL3* and *ST3GAL6* were significantly downregulated by both DHT and R1881. Of note, *ST6GAL1* which has previously been reported to be positively regulated by androgens did not increase with either DHT or R1881^28^. Similarly, transcriptomic analysis of matched patient biopsies revealed that in response to androgen deprivation therapy, *ST3GAL1* was significantly upregulated, and *ST6GALNAC1* was downregulated (**figure 4b**)^66^. *ST6GAL1* remained unchanged following androgen deprivation therapy. These findings were confirmed by treating LNCaP and VCaP cells with the second-generation AR-antagonists darolutamide and enzalutamide. In response to AR-inhibition, ST3Gal1 was significantly upregulated at both the gene and protein level (**figure 4c-e and supplemental figure 3a-c**), and ST6GalNAc1 was repressed. ST6Gal1 remained unchanged in response to AR-inhibition. We have previously confirmed that α2-3-linked sialic acid, added by ST3Gal1 is upregulated in response to AR-antagonism^30^. Flow cytometry with the SNA lectin was used to probe α2-6-sialic acid on the surface of LNCaP cells. Treatment with 1 nM R1881 (**figure 4f**) and the AR-inhibitor darolutamide (**figure 4g**) resulted in no change in α2-6-sialic acid, indicating that α2-6-linked sialoglycans typically synthesised by ST6Gal1 are not regulated by androgens. The sialyl-Tn (sTn) antigen has been robustly shown to be made by ST6Gal1Nac1, and shown to be AR-regulated in LNCaP cells by immunofluorescence^29^. In line with findings on breast cancer cell lines from Murugesan *et al*, we have shown LNCaP cells to have very low levels of sTn expression detected by flow cytometry (**supplemental figure 3d**)^67^. Using the K562 leukaemia cell line as a positive control, we found that the LNCaP, PC3 and PNT1A cell lines have little sTn expression as determined by flow cytometry (**supplemental figure 3d**). There was no detectable induction of sTn expression following AR stimulation or antagonism as determined by immunofluorescence, however this may be due to very low levels of expression (**supplemental figure 3e-f**). As such, we have been unable to confirm previous findings that the sTn antigen is indeed AR-regulated in LNCaP cells^26^.

**Figure 4.**
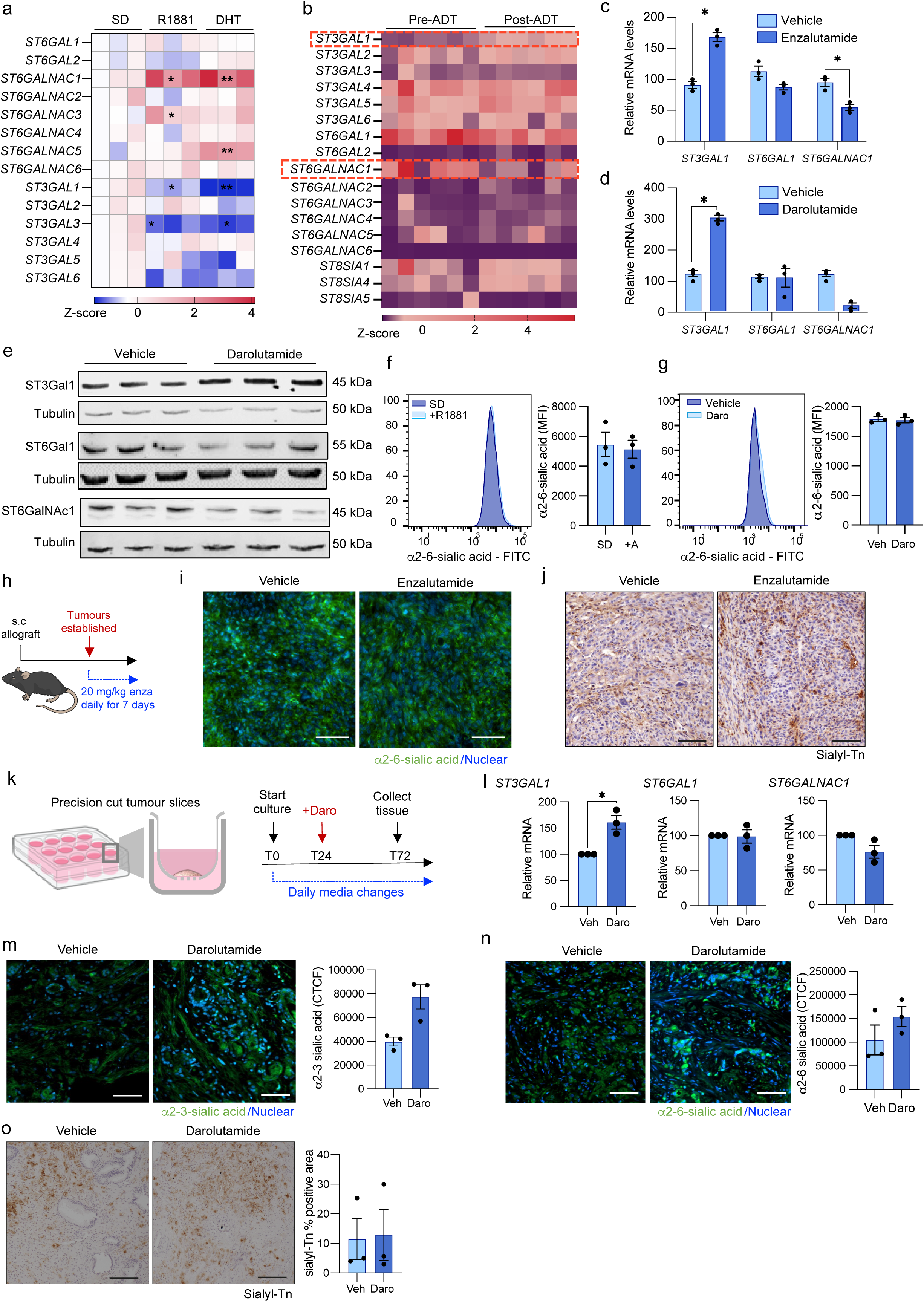
Sialyltransferases with preferences for *O*-glycans are modulated by androgens. (**a**) Heatmap showing gene expression as determined by quantitative-PCR, of a panel of sialyltransferases across serum deprived (SD), 1 nM R1881 and 10 nM DHT treatment. N = 3 biologically independent samples. Statistical significance determined using a one-way ANOVA. (**b**) Heatmap showing gene expression of sialyltransferases in transcriptomic data from matched patient biopsies pre-ADT and post-ADT. N = 8. Dataset from Long *et al.*^66^. (**c-d**) *ST3GAL1*, *ST6GAL1* and *ST6GALNAC1* mRNA expression in LNCaP cells following treatment with 10 µM enzalutamide or 10 µM darolutamide (compred with DMSO control) for 48 hours measured by RT-qPCR. N = 3 biologically independent replicates. Statistical significance determined using a paired two-way *t*-test. (**e**) Western blot analysis of ST3Gal1, ST6Gal1 and ST6GalNAc1 expression post treatment of LNCaP cells with 10 µM darolutamide (or DMSO control) for 48 hours. N = 3 biological replicates. (**f**) SNA lectin flow cytometry detection of α2-6 sialic acid on the surface of LNCaP cells following treatment with 1 nM synthetic androgen R1881 (or steroid depleted) for 24 hours by flow cytometry. Representative histogram of N =3 biologically independent samples shown. Statistical significance determined using a paired two-way *t*-test. (**g**) SNA lectin flow cytometry detection of α2-6 sialic acid on the surface of LNCaP cells following treatment with DMSO vehicle control (vehicle) and 10 µM darolutamide (Daro) for 48 hours. Representative histogram of N =3 biologically independent samples shown. Statistical significance determined using a paired two-way *t*-test. (**h**) Schematic of subcutaneous mouse model. 2 x 10^6^ TRAMP-C2 cells were implanted into the flank of C57BL/6 mice. Once tumours were established mice were given a daily dose of 20 mg/kg enzalutamide by oral gavage. NIAID Visual & Medical Arts. (10/7/2024). Lab Mouse. NIAID NIH BIOART Source. bioart.niaid.nih.gov/bioart/279. (**i**) Representative images of immunofluorescent staining of SNA which detects α2-6 sialic acid in mouse allograft tumours following treatment with vehicle control (N = 5) and enzalutamide (N = 6). Quantification of cellular total cell fluorescence of α2-6 sialic acid expression between mouse allograft tumours treated with enzalutamide (N = 6) and vehicle control (N = 5). Scale bar = 100 µm. (**j**) Representative images of immunohistochemical staining of sialyl-Tn in mouse allograft tumours following treatment with vehicle control (N = 5) and enzalutamide (N = 6). (**k**) Schematic oh *ex vivo* precision cut tumour slices treated with darolutamide. Tumour cores were precision cut to a thickness of 200 µm. Slices were allowed to recover for 24 hours and treated with either DMSO or darolutamide (10 µM) for 48 hours. NIAID Visual & Medical Arts. (10/7/2024). 12 Well Plate. NIAID NIH BIOART Source. bioart.niaid.nih.gov/bioart/3. (**l**) Gene expression of *ST3GAL1, ST6GAL1* and *ST6GALNAC1* in treated precision cut tumour slices as determined by q-PCR. N = 3 biologically independent replicates. Statistical significance determined using a paired two-way t-test. (**m**) Representative images of immunofluorescent staining of MAL-II which detects α2-3 sialic acid in precision cut tumour slices following treatment with vehicle control (N = 3) and darolutamide (N = 3). Quantification of cellular total cell fluorescence of α2-3 sialic acid expression between groups. N = 3 biologically independent replicates. Statistical significance determined using a paired two-way *t*-test. (**n**) Representative images of immunofluorescent staining of SNA which detects α2-6 sialic acid in precision cut tumour slices following treatment with vehicle control (N = 3) and darolutamide (N = 3). Quantification of cellular total cell fluorescence of α2-6 sialic acid expression between groups. N = 3 biologically independent replicates. Statistical significance determined using a paired two-way *t*-test. (**o**) Representative images of immunohistochemical staining of sialyl-Tn in precision cut tumour slices following treatment with vehicle control (N = 3) and darolutamide (N = 3). Quantification of the percentage of positive area shown. Data points shown are the mean of quantification of 3 regions of interest. N = 3 biologically independent replicates. Statistical significance determined using a paired two-way *t*-test.

To assess AR modulation of α2-6-sialic acid and the sTn antigen *in vivo*, we dosed C57BL/6 mice harbouring TRAMP-C2 subcutaneous allografts with the AR-antagonist enzalutamide daily for 7 days (**figure 4h**). Immunofluorescent and immunohistochemical staining confirmed that AR-targeting does not alter α2-6-linked sialic acid or the sTn antigen *in vivo* (**figure 4i-j and supplemental figure 4g-4h**). We noted that although some positive sTn antigen staining was detected in tumour cells within the allografts, we noted highest expression in stromal regions. We sought to confirm these findings in a human *ex vivo* precision cut tumour slice model. Tumour tissue was surgically removed from patients, precision sliced with a vibratome and cultured in transwells as described previously^42^. Tumour slices were rested for 24 hours and then treated with 10 µM darolutamide or vehicle control for 48 hours (**figure 4k**). Treatment with the AR-antagonist resulted in a significant reduction in expression of the AR-target gene *KLK3* (**supplemental figure 3i**). In tumour slices treated with darolutamide there was a significant increase in gene expression of *ST3GAL1* and a small non-significant decrease in *ST6GALNAC1*. Levels of *ST6GAL1* remained unchanged (**figure 4l**). Immunofluorescent staining confirmed that AR-targeting with darolutamide is associated with an upregulation α2-3-linked sialic acid. Whilst no change in the in expression of α2-6-linked sialic acid or sTn antigen was observed (**figure 4m-o**).

Using a wide range of strategies and models, this work confirms that ST3Gal1 and ST6GalNAc1 are altered in response to AR signalling. Our data confirms that α2-3-sialic acid, like ST3Gal1, is repressed by the AR and that ST6Gal1 and the glycans it synthesises are not AR-regulated.

### Supraphysiological levels of androgens induce differential ST patterns in prostate cancer

Previously, Munkley *et al.* reported that ST6Gal1 and α2-6-sialic acid was induced by androgens, in contrast with our findings presented here^28^. We noted that this study used 10 nM R1881, commonly used to model supraphysiological androgen levels (SPA). Bipolar androgen therapy (BAT) is a relatively new approach, whereby patients are cycled through SPA and near-castrate levels of androgens to resensitise them to conventional hormone therapies. Downregulation of the c-MYC oncogene in response to SPA has been identified as a key therapeutic mechanism in BAT^68^. Treatment of LNCaP cells with 1 nM and 10 nM R1881 confirmed that c-MYC is downregulated in response to AR stimulation, with the biggest downregulation in response to SPA (**figure 5a**). Levels of the full-length androgen receptor remained unchanged (**supplemental figure 4a**). In line with our previous work, ST3Gal1 was downregulated in response to 1 nM R1881, however levels are comparable to steroid depleted levels with 10 nM R1881 (**figure 5a**). In contrast, ST6Gal1 levels remain unchanged with 1 nM R1881 but are upregulated following SPA, in line with previous findings^28^. Additionally, ST6Gal1NAc1 is upregulated following 1 nM R1881, but is downregulated by 10 nM of androgens. We found that treatment of LNCaP cells with 10 nM R1881 failed to induce changes in either α2-3- or α2-6-linked sialic acid levels (**figure 5b-c**). Glycopeptide profiling of LNCaP cells confirmed that expression profiles of glycopeptides (**figure 5d**) and sialylated glycopeptides (**figure 5e**) differ between cells treated with 1 nM and 10 nM of synthetic androgens. Following treatment with physiological androgens we detected more glycopeptides in LNCaP cells (f**igure 5f**). The number of glycopeptides identified in cells treated with SPA were comparable to steroid depleted controls. When we performed Hallmark analysis of proteins which are sialylated in response to androgen stimulation, we found that three ‘Hallmarks’ were significantly enriched, including Myc target genes (**figure 5g**). These data suggest that SPA conditions produce distinct glycoprotein profiles from physiological levels and that these differences are mediated through altered expression of STs at these concentrations.

**Figure 5.**
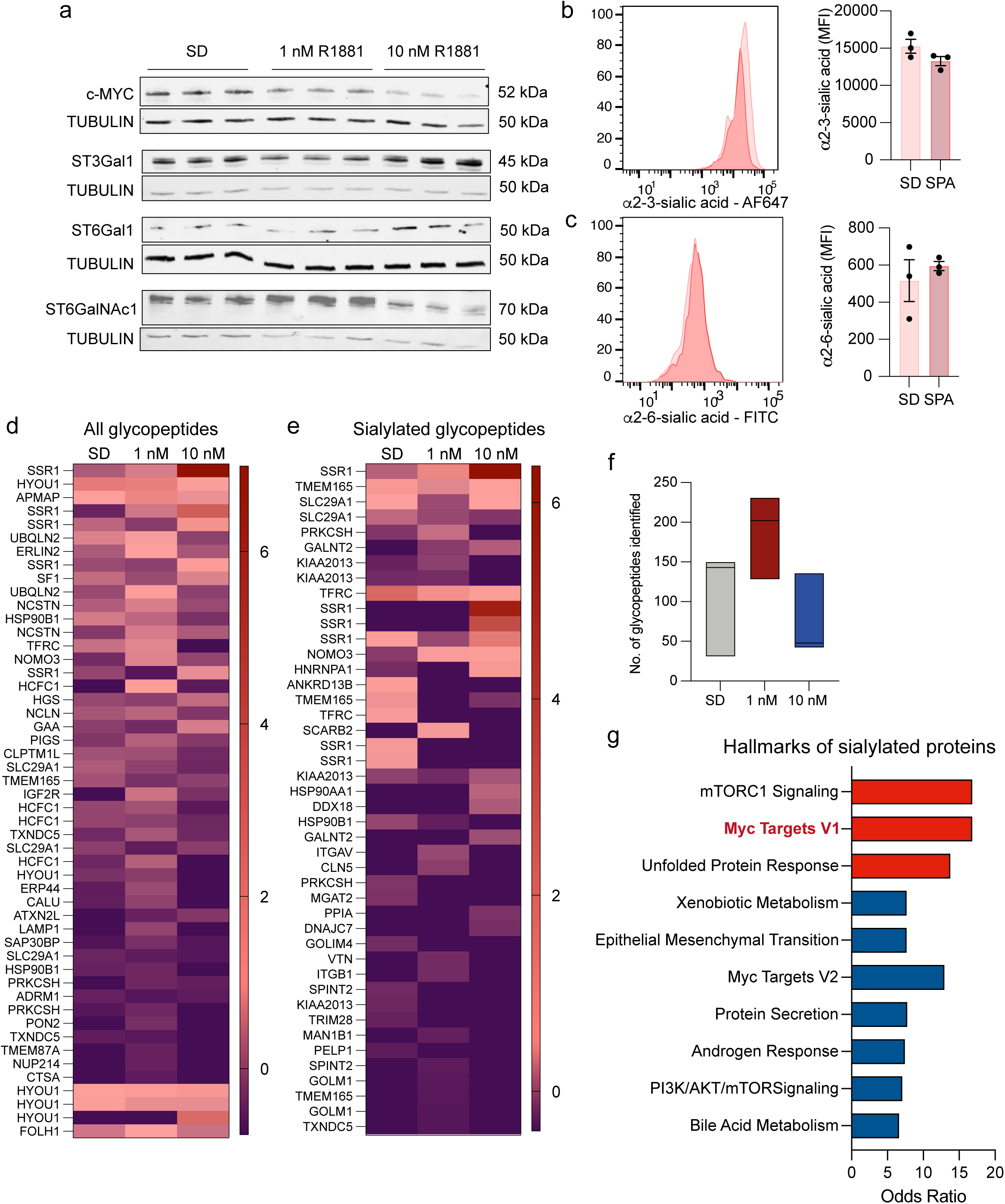
Supraphysiological levels of androgens induce differential expression of sialoglycans. (**a**) Western blot analysis of c-MYC, ST3Gal1, ST6Gal1 and ST6GalNAc1 expression of 72 hours steroid depleted LNCaP cells stimulated with 1 nM or 10 nM R1881 (synthetic androgen) for 24 hours. Representative image of N=3 biological replicates. (**b**) MAL-II lectin detection of α2-3 sialic acid in 72 hours steroid depleted LNCaP cells following 24 hours treatment with SPA (10 nM R1881) by flow cytometry. Representative histogram and bar chart of median fluorescent intensities shown. Histogram representative of N = 3 biologically independent samples. Statistical significance determined using a paired two-way *t*-test. (**c**) SNA lectin detection of α2-6 sialic acid in 72 hours steroid depleted LNCaP cells following 24 hours treatment with SPA (10 nM R1881) by flow cytometry. Representative histogram and bar chart of median fluorescent intensities shown. Histogram representative of N = 3 biologically independent samples. Statistical significance determined using a paired two-way *t*-test. (**d**) Heatmap showing z-scores of differentially glycosylated proteins in LNCaP cells under steroid depleted conditions or treated with 1 nM or 10 nM R1881 for 24 hours. N= 3, each cell is the mean of 3 technical replicates. Z-scores calculated from the relative MS intensity. (**e**) Heatmap showing z-scores of differentially sialylated proteins in LNCaP cells under steroid depleted conditions or treated with 1 nM or 10 nM R1881 for 24 hours. N= 3, each cell is the mean of 3 technical replicates. Z-scores calculated from the relative MS intensity. (**f**) Box plot of number of unique glycopeptides identified in LNCaP cells under steroid depleted conditions or treated with 1 nM or 10 nM R1881 for 24 hours, N=3. (**g**) Bar chart showing Hallmarks that are enriched in proteins which are differentially sialylated in R1881 treated LNCaP cells. Significantly enriched Hallmarks are highlighted with red bars. Hallmark analysis was performed using Enrichr (https://maayanlab.cloud/Enrichr/).

### c-MYC and the AR bind to regulatory regions in ST3Gal1 to mediate α2-3-linked sialylation in prostate cancer

Having observed differential expression of STs following treatment with SPA where c-MYC is repressed, we next sought to confirm whether the c-MYC itself can modulate expression of STs in prostate cancer. c-MYC knockdown resulted in increased gene expression of *ST3GAL1* (**figure 6a**) and expression of the ST3Gal1 enzyme (**figure 6b**). Neither *ST6GAL1* nor *ST6GALNAC1* were altered following loss of MYC. Similarly, loss of MYC resulted in an increase in expression of α2-3-linked sialic acid (**figure 6c**) and a concurrent decrease in levels in unsialylated galactose as shown by binding of Peanut Agglutinin (PNA) lectin (**figure 6d**). Levels of α2-6-linked sialic acid remained unchanged (**supplemental figure 5a**). We next asked whether the same effect could be seen in prostate cancer patients. MYC transcriptomic activity signatures can be better prognostic markers than MYC-amplification status alone, and as such we used the MYC activity score described by *Schaafsma et al*. (2020) to assess the effect of MYC activity on ST expression in The Cancer Genome Atlas (TCGA) prostate adenocarcinoma (PRAD) cohort^69^. This analysis confirmed that in primary prostate cancer patients, high MYC activity is associated with a significant reduction in *ST3GAL1* (**figure 6e**). A reduction in *ST6GAL1* expression was also associated with high MYC activity, whereas no relationship was observed between *ST6GALNAC1* and MYC activity. We next used iPSCs with engineered PTEN- and TP53-loss to investigate the effect of MYC overexpression on ST3Gal1-synthesised sialoglycans in prostate cancer organoids. Prostate cancer organoids were generated as described in Shaikh et al. (under review). Organoids with a MYC overexpression were compared with PTEN- and TP53 null controls and showed a reduction in α2-3-linked sialoglycans (**figure 6f**).

**Figure 6.**
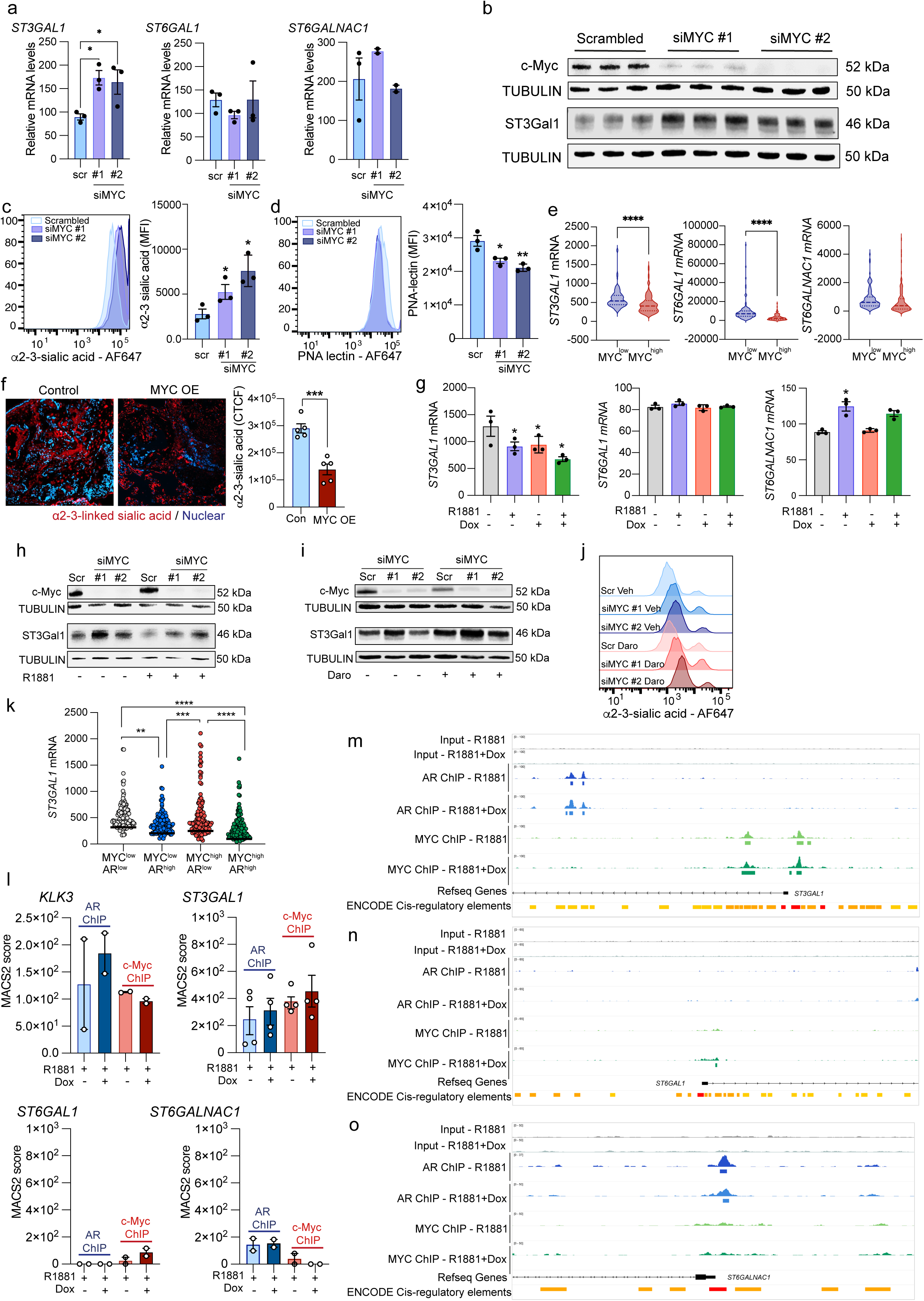
MYC and the AR cooperate to control ST3Gal1-mediated sialylation in prostate cancer. (**a**) qPCR analysis of *ST3GAL1, ST6GAL1* and *ST6GALNAC1* gene expression in LNCaP cells following gene targeting with scrambled siRNA, siMYC #1 and siMYC #2 treatment for 48 hours. Statistical significance determined using a one-way ANOVA. (**b**) Western blot analysis of c-MYC and ST3Gal1 expression of in LNCaP cells treated with scrambled siRNA, siMYC #1 or siMYC #2. Representative image of N=3 biological replicates. (**c**) MAL-II lectin detection of α2-3 sialic acid in LNCaP cells following treatment with scrambled siRNA, siMYC #1 and siMYC #2 treatment for 48 hours by flow cytometry. Representative histogram and bar chart of median fluorescent intensities shown. N = 3 biologically independent samples. Statistical significance determined using a one-way ANOVA. (**d**) PNA lectin detection of desialylated galactose in LNCaP cells following treatment with scrambled siRNA, siMYC #1 and siMYC #2 treatment for 48 hours by flow cytometry. Representative histogram and bar chart of median fluorescent intensities shown. N = 3 biologically independent samples. Statistical significance determined using a one-way ANOVA. (**e**) Transcriptomic profiling of *ST3GAL1, ST6GAL1* and *ST6GALNAC1* in patients in the prostate adenocarcinoma (PRAD) The Cancer Genome Atlas (TCGA) cohort stratified by MYC activity status (N = 245). Statistical significance determined using a one-way ANOVA. (**f**) Representative images of immunofluorescent staining of MAL-II which detects α2-3-sialic acid in prostate cancer organoids with c-MYC overexpression. Quantification of cellular total cell fluorescence of α2-3-sialic acid expression in control and c-MYC overexpressing organoids. (**g**) Transcriptomic profiling of *ST3GAL1*, *ST6GAL1* and *ST6GALNAC1* in LNCaP treated with 1 nM R1881 and/or 2 mg/mL doxycycline to induce MYC overexpression. N = 3 biological replicates (GEO: GSE73917) ^15^. (**h**) Western blot analysis of c-MYC and ST3Gal1 expression of in LNCaP cells treated with scrambled siRNA, siMYC #1 or siMYC #2 in steroid depleted conditions ± 1 nM R1881. Representative image of N=3 biological replicates. (**i**) Western blot analysis of c-MYC and ST3Gal1 expression in LNCaP cells treated with scrambled siRNA, siMYC #1 or siMYC #2 ± 10 µM darolutamide. Representative image of N=3 biological replicates. (**j**) MAL-II lectin detection of α2-3 sialic acid in LNCaP cells following treatment with scrambled siRNA, siMYC #1 and siMYC #2 ± 10 µM darolutamide by flow cytometry. Representative histogram and bar chart of median fluorescent intensities shown. N = 3 biologically independent samples. Statistical significance determined using a two-way ANOVA. (**k**) Transcriptomic profiling of *ST3GAL1, in* patients in the prostate adenocarcinoma (PRAD) The Cancer Genome Atlas (TCGA) cohort stratified by MYC and AR activity status (N = 245). Statistical significance determined using a one-way ANOVA. (**l**) ChIP-seq signal for AR and MYC in either the presence or absence of MYC overexpression (+Dox) over the promoter region of the indicated genes as calculated by MACS2. Data from GSE73994^15^. (**m-o**) Snapshot from the IGV genome browser over the promoter regions of ST3GAL1, ST6GAL1 and ST6GALNAC1 respectively, showing AR and MYC ChIP-seq coverage in either the presence or absence of MYC overexpression (+Dox). Bars at the bottom of each track indicate peaks identified by MACS2. Input data is shown as control. ENCODE cis-regulatory regions are shown following the UCSC display conventions, e.g. red indicates promoter-like signature, orange indicates proximal enhancer-like signature and yellow indicates distal enhancer-like signature Data from GSE73994 ^15^.

Given that we have shown both AR- and MYC-dependent changes in prostate tumour sialoglycan composition, we next sought to understand the combined effect of the two transcription factors on ST expression. Using LNCaP cells with doxycycline inducible MYC overexpression treated with 1 nM R1881 it was reported that the AR and MYC share 25-30% of their target binding sites^15^. Barfeld *et al.* discovered that 25% of shared AR-MYC targets have an antagonistic relationship, with the two transcription factors working together to positively regulate 1.5% of target genes. Analysis of previously published data showed that androgen stimulation and MYC overexpression alone both resulted in significant reductions in *ST3GAL1* expression compared with controls^15^. Importantly, treatment with androgens in combination with MYC overexpression resulted in the largest reduction in *ST3GAL1* transcripts. Neither stimulation with R1881 or MYC-overexpression altered *ST6GAL1* expression. In line with our previous findings, androgen stimulation significantly increased *ST6GALNAC1* expression and modulation of the MYC oncogene alone had no effect on *ST6GALNAC1* expression. (**figure 6g**). We confirmed this effect in LNCaP cells with siRNA knockdown of c-MYC and treatment with 1 nM of R1881. As we have shown previously, loss of c-MYC in steroid depleted conditions led to a significant increase in ST3Gal1 (**figure 6h**). In cells treated with synthetic androgens, siRNA knockdown of MYC also led to an increase in ST3Gal1, however to a lesser extent (**figure 6h**). Whilst, in cells treated with darolutamide, inhibition of the androgen signalling cascade cooperates with loss of c-MYC to upregulate the expression of ST3Gal1 and the expression of α2-3-linked sialoglycans to a greater extent than c-MYC alone (**figure 6i-j**). These data suggest that the two transcription factors may synergise to repress expression of *ST3GAL1*. We confirmed that the same effect is observed in prostate cancer patients by stratifying patients in the TCGA PRAD cohort based on MYC and AR activity (N= 250), showing lowest ST3GAL1 expression in MYC^high^/AR^high^ patients (**figure 6k**).

We next analysed previously published AR and c-MYC ChIP-sequencing data from LNCaP cells with doxycycline inducible MYC overexpression treated with 1 nM R1881 (GSE73994)^13^ Model-based Analysis of ChIP-seq (MACS2) scores suggest that both the AR and MYC bind to regulatory elements close to the promoter of *ST3GAL1* (*KLK3* was used a positive control) (**figure 6l and supplemental figure 5b-5c**). Overexpression of MYC does not alter AR binding (**figure 6m**). We observe no binding of the AR following treatment with R1881 to regulatory regions in ST6Gal1, and some binding of MYC following MYC overexpression (**figure 6l, 6n and supplemental figure 5d**). Following treatment with R1881, the AR binds to regulatory regions in ST6GalNAc1, and we observe no binding of c-MYC (**figure 6l, 5o and supplemental figure 5e**). This confirms a cooperative relationship between the AR and MYC and suggest for the first time that both transcription factors directly bind to regulatory regions in *ST3GAL1* which may repress its expression.

### Synergy between the AR and MYC represses immunosuppressive Siglec-7 ligands

Given that SPA induces differential expression of sialoglycans it is reasonable to hypothesise that differential levels of androgen exposure may have differing consequences for immunosuppressive Siglec ligands. Previously, we have reported that ST3Gal1 synthesises Siglec-7 and Siglec-9 ligands in an androgen-dependent manner. As we have shown here that Siglec-7/-9 and -15 are highly expressed on PCa cells, we assessed the effect of normal physiological androgen exposure, and SPA on their expression on the PCa cell surface. Consistent with previous findings, Siglec-7 are reduced with normal androgen stimulation (**figure 7a-b**). We observed no difference in Siglec-7/-9 ligand expression between steroid depleted controls and 10 nM R1881. We observed no difference in Siglec-15 ligand expression with 1 nM R1881, and a minimal reduction in their expression with 10 nM R1881 (**figure 7c**). As we see the biggest change in Siglec-7 ligands in response to AR modulation, we sought to confirm that changes in Siglec ligand patterns in response to different concentrations of androgens are mediated through ST3Gal1. We utilised a previously characterised LNCaP shST3Gal1 knockdown cell line which has decreased cell surface α2-3-sialoglycans to investigate the effects of normal SPA stimulation (**supplemental figure 6a and 6b**)^30^. In control cells, exposure to normal and SPA conditions resulted in distinct Siglec-7 ligand expression profiles. This differential response was abrogated in ST3Gal1-deficient LNCaP cells, indicating that ST3Gal1 is required for androgen-mediated modulation of Siglec-7 ligands (**figure 7d**). To further validate our findings, by mutating histidine 299 to an alanine residue, and expressing this in LNCaP-shST3Gal1 cells we generated cells which express a catalytic-dead version of ST3Gal1 (LNCaP-ST3Gal1^H299A^)^70^. We confirmed that ST3Gal1^H299A^ cells express the same level of ST3Gal1 as WT control cells (**figure 7e**) but display a reduction in Siglec-7 ligands on their cell surface compared with ST3Gal1^WT^ controls (**figure 7f**). As with LNCaP-shST3Gal1 cells, in LNCaP-ST3Gal1^H299A^ cells physiological and SPA fail to induce to significant differences in both α2-3-sialoglycans (**supplemental figure 6c**) and Siglec-7 ligands on PCa cells (**figure 7g**). These data confirm that ST3Gal1 is a critical ST responsible for mediating dynamic AR-responsive changes in Siglec-7 ligand expression.

**Figure 7.**
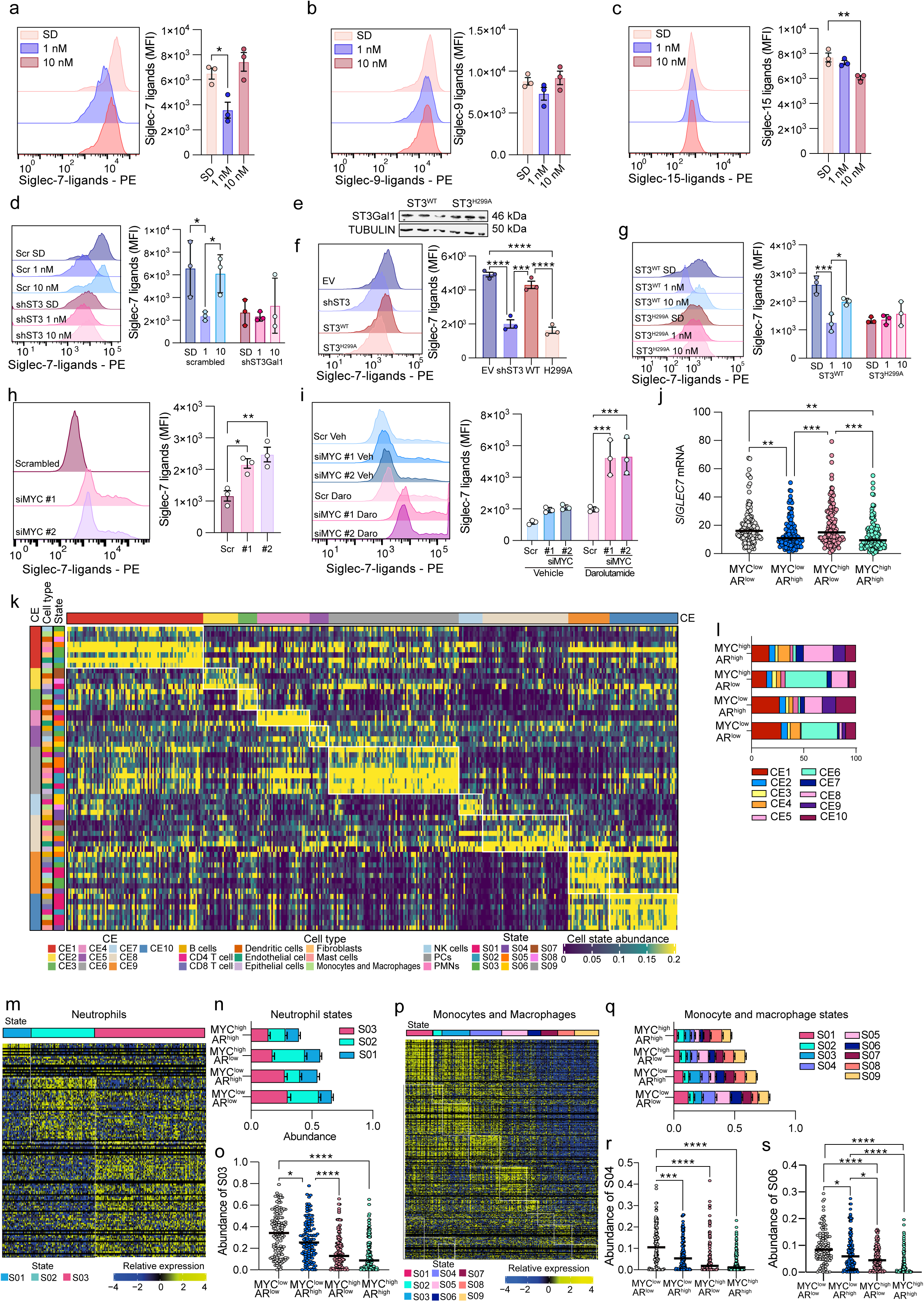
The AR and MYC work together to regulate the synthesis of Siglec-7 ligands in prostate cancer. (**a-c**) Siglec-7, Siglec-9, and Siglec-15 ligand detection in LNCaP cells steroid depleted for 72 hours and then stimulated with either 1 nM or 10 nM R1881 for 24 hours and quantified by flow cytometry. Representative histogram and bar chart of median fluorescent intensities shown. Histogram and bar chart representative of N = 3 biologically independent samples. Statistical significance determined using a one-way ANOVA. (**d**) Siglec-7 ligand detection across scrambled control and shST3 LNCaP cells steroid depleted for 72 hours and stimulated with either 1 nM or 10 nM R1881 for 24 hours by flow cytometry. Representative histogram and bar chart of median fluorescent intensities shown. Histogram and bar chart representative of N = 3 biologically independent samples. Statistical significance determined using a two-way ANOVA. (**e**) Western blot analysis of ST3Gal1 expression of in shST3Gal1-LNCaP cells transfected with wildtype ST3Gal1 (ST3Gal1^WT^) or catalytic-dead ST3Gal1 (ST3Gal1^H299A^) (**f**) Siglec-7 ligand detection across scrambled control and shST3 LNCaP cells and shST3Gal1 cells transfected with wildtype (WT) ST3Gal1 or catalytic dead ST3Gal1 (H299A) by flow cytometry. Representative histogram and bar chart of median fluorescent intensities shown. Histogram and bar chart representative of N = 3 biologically independent samples. Statistical significance determined using a two-way ANOVA. (**g**) Siglec-7 ligand detection in shST3Gal1 cells transfected with wildtype ST3Gal1 (ST3Gal1^WT^) or catalytic-dead ST3Gal1 (ST3Gal1^H299A^) by flow cytometry. Representative histogram and bar chart of median fluorescent intensities shown. Histogram and bar chart representative of N = 3 biologically independent samples. Statistical significance determined using a two-way ANOVA. (**h**) Siglec-7 ligand detection in LNCaP cells treated with scrambled siRNA or two different siRNA’s targeting c-MYC for 48 hours quantified by flow cytometry. Representative histogram and bar chart of median fluorescent intensities shown. Histogram and bar chart representative of N = 3 biologically independent samples. Statistical significance determined using a two-way ANOVA. (**i**) Siglec-7 ligand detection in LNCaP cells treated with scrambled siRNA or two different siRNA’s targeting c-MYC ± 10 µM darolutamide quantified by flow cytometry. Representative histogram and bar chart of median fluorescent intensities shown. Histogram and bar chart representative of N = 3 biologically independent samples. Statistical significance determined using a two-way ANOVA. (**j**) Transcriptomic profiling of *SIGLEC7, in* patients in the prostate adenocarcinoma (PRAD) The Cancer Genome Atlas (TCGA) cohort stratified by MYC and AR activity status (N = 245). Statistical significance determined using a one-way ANOVA. (**k**) Heatmap summarising the Ecotypes (CE), Cell Types, and Cell States identified in patients in the prostate adenocarcinoma (PRAD) The Cancer Genome Atlas (TCGA) cohort. Analysis run using Ecotyper (https://ecotyper.stanford.edu/) ^39^. (**l**) Stacked bar chart showing the percentage of patients in the prostate adenocarcinoma (PRAD) The Cancer Genome Atlas (TCGA) cohort assigned to each of 10 cellular ecotypes (CE) with patients stratified by MYC and AR activity. (**m**) Heatmap summarising the three neutrophil states identified by Ecotyper in the PRAD TCGA cohort. (**n**) Stacked bar chart of the mean of the absolute values of each neutrophil state in patients stratified by MYC and AR activity. (**o**) Data shows the abundance of cell state 3 (S03) neutrophils in patients stratified by MYC and AR activity. Statistical significance determined using a one-way ANOVA. (**p**) Heatmap summarising the nine monocyte/macrophage states identified by Ecotyper in the PRAD TCGA cohort. (**q**) Stacked bar chart of the mean of the absolute values of each monocyte/macrophage state in patients stratified by MYC and AR activity. (**r**) Data shows the abundance of cell state 4 (S04) macrophages in patients stratified by MYC and AR activity. Statistical significance determined using a one-way ANOVA. (**s**) Data shows the abundance of cell state 6 (S06) macrophages in patients stratified by MYC and AR activity. Statistical significance determined using a one-way ANOVA.

As we have now shown ST3Gal1 is modulated by both the AR and MYC in response to SPA and have found distinct Siglec-7 ligands in response to differing concentrations of androgens, we next asked how c-MYC may contribute to Siglec-7 ligand expression in PCa. Previous studies have reported that c-MYC positively regulated Siglec-7 ligands through ST6GalNAc4 in leukaemia^36^. Following gene knockdown of MYC, Siglec-7 ligands were significantly increased in LNCaP cells (**figure 7h**). We then explored the combined effect of gene knockdown of MYC with AR antagonism following Darolutamide treatment on Siglec-7 ligands. This revealed that both MYC knockdown and AR antagonism coordinate to upregulate the expression Siglec-7 ligands. We sought to explore these findings in prostate cancer patients. We stratified patients in the TCGA-PRAD (N = 250) cohort by MYC and AR activity and found that mRNA expression of *SIGLEC7* is significantly reduced in MYC^high^/AR^high^ patients (**figure 7j**). As *SIGLEC7* is an immunoreceptor, this data suggests that MYC^high^/AR^high^ patients may have a reduced number of Siglec-7^+^ myeloid cells.

As our data suggests that the Siglec-7 receptor and its ligands are repressed by the AR and MYC, we hypothesised that increased MYC and AR activity may remove myeloid-driven immune suppression in prostate cancer. To this end, we utilised Ecotyper, a machine learning framework which allows for identification of cell subsets or ‘states’ and multicellular niches or ‘ecotypes’ from bulk-transcriptomic profiling^39^. We performed Ecotyper analysis on data available in the TCGA PRAD dataset and were able to assign Ecotypes to 385 patients (**figure 7k**). We then stratified patients based on AR and MYC activity as we had done previously, and identified ecotypes associated with AR and MYC activity (**figure 7l**). We identified significant differences in ecotype composition in AR^high^/MYC^high^ patients. Significant changes included a decrease in ecotype 1, associated with few infiltrating immune cells and poor clinical outcomes. The abundance of ecotype 9 was 229% higher in MYC^high^/AR^high^ patients compared with MYC^low^/AR^low^ patients. Ecotype 9 has been shown to have significant power in predicting patient response to immune checkpoint inhibitors (ICIs), suggesting that AR^high^/MYC^high^ patients may have clinically favourable responses to ICIs^39^.

As we had previously shown Siglec-7 to be expressed on macrophages and neutrophils and given the role of ST3Gal1 and Siglec-7 in myeloid-mediated immune suppression in prostate tumours we interrogated the cell states of these myeloid cells in MYC^high^/AR^high^ patients. Three subsets of neutrophils were identified based on cell states (**figure 7m**). When comparing neutrophil subsets between patient groups, we observed a stepwise reduction in state 3 neutrophils (**figure 7n**). Neutrophils classed as state 3 by Ecotyper express genes typically associated with immunosuppressive pro-tumour neutrophils such as *MRC1* (CD206), *MMP8*, *ARG1* and *OLR1 (*LOX-1) ^71–77^. The number of these suppressive neutrophils was significantly decreased in MYC^high^/AR^high^ patients compared with MYC^low^/AR^low^ patients (**figure 7o**). Ecotyper identified 9 cellular states of monocytes and macrophages in the TCGA PRAD dataset (**figure 7p**). We observed MYC and/or AR dependant changes in macrophage states (**figure 7q**). We found that the largest changes in macrophage numbers between patient subgroups were in macrophage states 4 and 6. State 4 and 6 macrophages are annotated by Ecotyper as ‘M2’ anti-inflammatory macrophage subsets with pro-tumour functions strongly associated with poor patient survival outcomes^39^. MYC^high^/AR^high^ patients have a ten-fold reduction in the abundance of state 4 macrophages and an eight-fold reduction in state 6 macrophages compared with MYC^low^/AR^low^ patients (**figure 7r and 7s**). Here, we conclude that the AR and MYC repress immunosuppressive Siglec-7 signalling, and that this may result in decreased suppressive myeloid populations in prostate tumours.

## Discussion

Tumour-associated glycosylation has emerged as a critical interface between oncogenic signalling and immune evasion^22,23,36,78^. Here, we uncover a synergistic mechanism by which the AR and the MYC oncogene regulate immunosuppressive sialoglycan biosynthesis in PCa. Specifically, we demonstrate that the AR and MYC transcription factors directly bind to and repress the α2,3-sialyltransferase ST3Gal1, leading to reduced expression of Siglec-7 ligands on the tumour cell surface.

Many studies have previously described an antagonistic relationship between the AR and MYC and several examples of this have been demonstrated^5,9,12,13,15–17,20,36,79^. A previous study found that a very small percentage of genes in the genome are repressed by both the AR and MYC and our findings identify *ST3GAL1* as one such example^15^. We show that both transcription factors bind to regulatory regions in the *ST3GAL1* gene and independently repress its expression, whilst their combined activity leads to synergistic suppression. In line with previous reports, MYC overexpression does not affect binding of the AR, possibly suggesting that the AR is the dominant transcription factor^15^. Additionally, we show that androgen concentrations significantly influence ST expression, revealing a threshold-sensitive effect on glycosylation pathways. Our work contributes to a growing body of literature which suggests a broader and more context-dependent spectrum of AR-MYC crosstalk than previously appreciated, potentially involving co-occupancy at repressive chromatin regions or shared recruitment of corepressors.

Recent work has highlighted c-MYC as a key transcription factor which positively regulates ST6GalNAc4 and the sialoglycans which it synthesises, leading to an increase in Siglec-7 ligands in leukaemia^36^. This is juxtaposed with our findings that in prostate tumours c-MYC represses Siglec-7 ligands. These results contribute to an increasing number of studies which reflect a tissue-specific regulatory architecture of sialoglycans and glyco-immune checkpoints, shaped by lineage-defining factors such as the AR. This highlights the need to interpret transcription factor dependant changes in the glycome within specific cellular contexts.

The repression of ST3Gal1 by the AR and MYC impacts the immune landscape of prostate tumours. The enzymatic activity of ST3Gal1 is responsible for generating Siglec-7 ligands, which are recognised by inhibitory receptors on tumour infiltrating neutrophils and tumour-associated macrophages. Our comprehensive analysis of cellular ecosystems in prostate cancer patients suggests that increases in AR and MYC signalling blunts this immunosuppressive signalling, decreasing populations of immunosuppressive myeloid subsets and generating an ecotype associated with pro-inflammatory signalling and improved responses to immune checkpoint inhibition. For the first time, we show that combined modulation of sialylation via hormonal and key oncogenic pathways may directly alter sialylation-mediated tumour immune evasion mechanisms in prostate cancer.

In summary, we present a previously unrecognised AR–MYC–ST3Gal1 axis that integrates oncogenic and hormonal transcriptional regulation with glyco-immune checkpoint control. These findings have implications for the use of androgen-based therapies in PCa and point to sialoglycans as actionable targets for immunomodulation.

## Acknowledgments

We thank CBC staff for the animal husbandry, Newcastle University Bioimaging unit staff for support with fluorescent imaging, Newcastle University Flow Cytometry Core Facility staff for assistance with flow cytometry. We thank Newcastle University Bioinformatics Support Unit for data analysis support and the University of York Bioscience Technology facility and Metabolomics and Proteomics Laboratory for assistance with glycopeptide analysis. K562 cells were a kind gift from Dr Lisa Russell (Newcastle University, UK). This work was funded by a Prostate Cancer UK Travelling Prize fellowship awarded to ES (TLD-PF19-002), a JGW Patterson Foundation grant awarded to ES and a Prostate Cancer UK Research Innovation Award awarded to ES (RIA23-ST2-008). The Newcastle Centre for Cancer is supported by Cancer Research UK Experimental Cancer Medicine Centre funding (25160). The York Centre of Excellence in Mass Spectrometry was created thanks to a major capital investment through Science City York, supported by Yorkshire Forward with funds from the Northern Way Initiative, and subsequent support from EPSRC (EP/K039660/1; EP/M028127/1). KC and ELW are supported by a Prostate Cancer Uk Research Innovation Award (RIA19-ST2-005). RM is supported by a Irish Cancer Society Translational Research Scholarship (CRS23MAG).

## Author contributions (CRediT statement)

**Emma Scott:** Conceptualisation, methodology, formal analysis, investigation, writing the original draft, visualisation, supervision, project administration and funding acquisition. **Adam Duxfield:** Conceptualisation, formal analysis, investigation, writing the original draft. **Rebecca Garnham:** Conceptualisation, formal analysis, investigation, writing the original draft. **Esme Hutton**: Formal analysis, investigation, review and editing of original draft. **Adam Dowle:** Data curation, formal analysis, review and editing of original draft. **Jamie Wills:** Formal analysis, investigation, review and editing of original draft. **Sara Luzzi:** Formal analysis, investigation, data curation, visualisation, review and editing of original draft. **Ryan Nelson:** Formal analysis, investigation, review and editing of original draft. **Emma Lishman-Walker:** Formal analysis, investigation, review and editing of original draft. **Rianna Magee:** Investigation, review and editing of original draft. **Adriana Buskin:** Investigation, review and editing of original draft. **Anastasia C. Hepburn:** Investigation, review and editing of original draft. **Tahsin Emirhan Tekoglu**: Formal analysis, investigation, visualisation, review and editing of original draft. **Kaidan Maloney-Friar**: Investigation, review and editing of original draft. **Fiona Frame:** Resources, review and editing of original draft. **Norman Maitland:** Resources, review and editing of original draft. **Ann Hedley:** Formal analysis, investigation, data curation, review and editing of original draft. **Holly Henderson**: Investigation, review and editing of original draft. **Benjamin McCullough:** Methodology, review and editing of original draft. **Bharat Gowardhan:** Resources, review and editing of original draft. **Kanagasabai Sahadevan:** Resources, review and editing of original draft. **Stuart McCracken:** Resources, review and editing of original draft. **Luke Gaughan:** Supervision, review and editing of original draft. **Rakesh Heer**: Supervision, review and editing of original draft. **Kelly Coffey:** Resources, supervision, review and editing of original draft. **Nathan Lack:** Supervision, review and editing of original draft. **Nathalie Signoret:** Supervision, review and editing of original draft. **Martin Fascione:** Supervision, review and editing of original draft.

## Figure legends

**Supplemental Figure 1.** mRNA levels of all twenty sialyltransferases in transcriptomic data from the SU2C PRAD cohort (Accessed through cBioPortal, N=208).

**Supplemental Figure 2.** (**a**) Schematic of flow cytometry gating strategies used for all MAL-II, SNA and Siglec-FC flow cytometry. (**b**) Bar chart of median fluorescent intensities shown for the quantification of the basal expression of CD22, CD33, Siglec-7, Siglec-9, Siglec-10 and Siglec-15 ligands on LNCaP cells, representative of N = 3 biologically independent samples. Data are mean ± standard error of the mean. (**c**) The CD22, CD33 and Siglec-10 immunoreceptors (red) co-localised with CD14 (green) in prostate cancer patient biopsies using dual immunofluorescence. Images were prepared using a ZEISS Axio Imager2 microscope with a x20 objective. Scale bar = 20 µm.

**Supplemental Figure 3.** (**a**) *ST3GAL1*, *ST6GAL1* and *ST6GALNAC1* mRNA expression in VCaP cells following treatment with vehicle control (DMSO) or 10 µM darolutamide for 72 hours measured by RT-qPCR. N = 3 biologically independent replicates. (**b**) Western blot analysis of androgen receptor (AR) expression of LNCaP cells treated with either vehicle control (DMSO) or 10 µM darolutamide for 72 hours. Representative image of N = 3 biological replicates. Loading control is shown by Tubulin expression (**c**) Western blot analysis of AR, ST3Gal1, ST6Gal1 and ST6GalNAc1 expression of LNCaP cells treated with either vehicle control (DMSO) or 10 µM enzalutamide for 72 hours. Representative image of N = 3 biological replicates. Loading control is shown by Tubulin expression. (**d**) Bar chart of median fluorescent intensities shown for the quantification of the basal expression of Sialyl T antigen (sTn) across K562, LNCaP, PC3 and PNT1A cell lines, representative of N = 3 biologically independent samples. (**e**) Representative images of immunofluorescent staining of sTn which detects the Sialyl-Tn antigen in LNCaP cells steroid depleted for 72 hours and stimulated with 1 nM R1881 (synthetic androgen) for 24 hours. (**f**) Representative images of immunofluorescent staining of sTn in LNCaP treated with vehicle control (DMSO) or 10 µM Darolutamide for 72 hours. (**g**) Representative images of immunofluorescent staining of SNA which detects α2-6 sialic acid in mouse allograft tumours following treatment with vehicle control (N = 5) and enzalutamide (N = 6). (**h**) Quantification of cellular total cell fluorescence of α2-6 sialic acid expression between mouse allograft tumours treated with enzalutamide (N = 6) and vehicle control (N = 5). (**i**) *KLK3* mRNA expression in LNCaP cells following treatment with vehicle control (DMSO) or 10 µM Darolutamide for 72 hours measured by RT-qPCR. N = 3 biologically independent replicates.

**Supplemental Figure 4.** Western blot analysis of androgen receptor (AR) expression of 72 hours steroid depleted LNCaP cells stimulated with 1 nM or 10 nM R1881 (synthetic androgen) for 24 hours. Representative image of N=3 biological replicates.

**Supplemental Figure 5.** (**a**) SNA lectin detection of α2-6 sialic acid in LNCaP cells following treatment with scrambled siRNA (scsiRNA) (N = 3), siMYC #1 (N = 3) and siMYC #2 (N = 3) for 48 hours by flow cytometry. Representative histogram and bar chart of median fluorescent intensities shown. N = 3 biologically independent samples. Statistical significance tested using a two-way ANOVA. Statistical significance is shown as * *p* < 0.05, ** (**b-e**) Snapshot from the IGV genome browser over the *KLK3* (PSA), *ST3GAL1*, *ST6GAL1* and *ST6GALNAC1* genes respectively, showing AR and MYC ChIP-seq coverage in either the presence or absence of MYC overexpression (+Dox). Bars at the bottom of each track indicate peaks identified by MACS2. ENCODE cis-regulatory regions are shown following the UCSC display conventions, e.g. red indicates promoter-like signature, orange indicates proximal enhancer-like signature and yellow indicates distal enhancer-like signature.

**Supplemental Figure 6.** (**a**) Western blot analysis of ST3Gal1 expression of scrambled control (Scr shRNA) and shST3 ST3Gal1 transduced LNCaP cells. Loading control is shown by Tubulin expression. (**b**) MAL-II lectin detection of α2-3 sialic acid in scrambled control (Scr shRNA) and shST3 ST3Gal1 transduced LNCaP cells by flow cytometry along with a secondary only control. Representative histogram and bar chart of median fluorescent intensities shown. Histogram and bar chart representative of N = 3 biologically independent samples. (**c**) MAL-II lectin detection of α2-3 sialic acid across scrambled control and shST3 LNCaP cells steroid depleted for 72 hours and stimulated with either 1 nM or 10 nM R1881 for 24 hours by flow cytometry. Representative histogram and bar chart of median fluorescent intensities shown. Histogram and bar chart representative of N = 3 biologically independent samples. Statistical significance determined using a two-way ANOVA.

