## Supplementary figures and images for "The Androgen Receptor and MYC synergise to modulate the synthesis of Siglec-7 ligands in prostate cancer"

### Supplemental figures

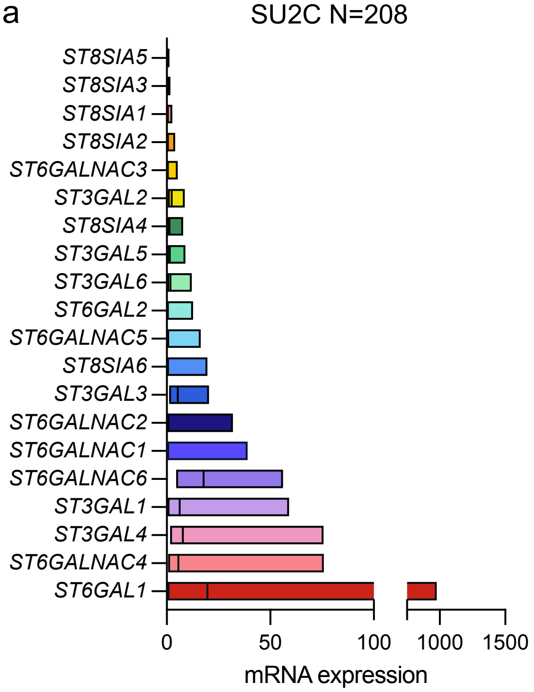

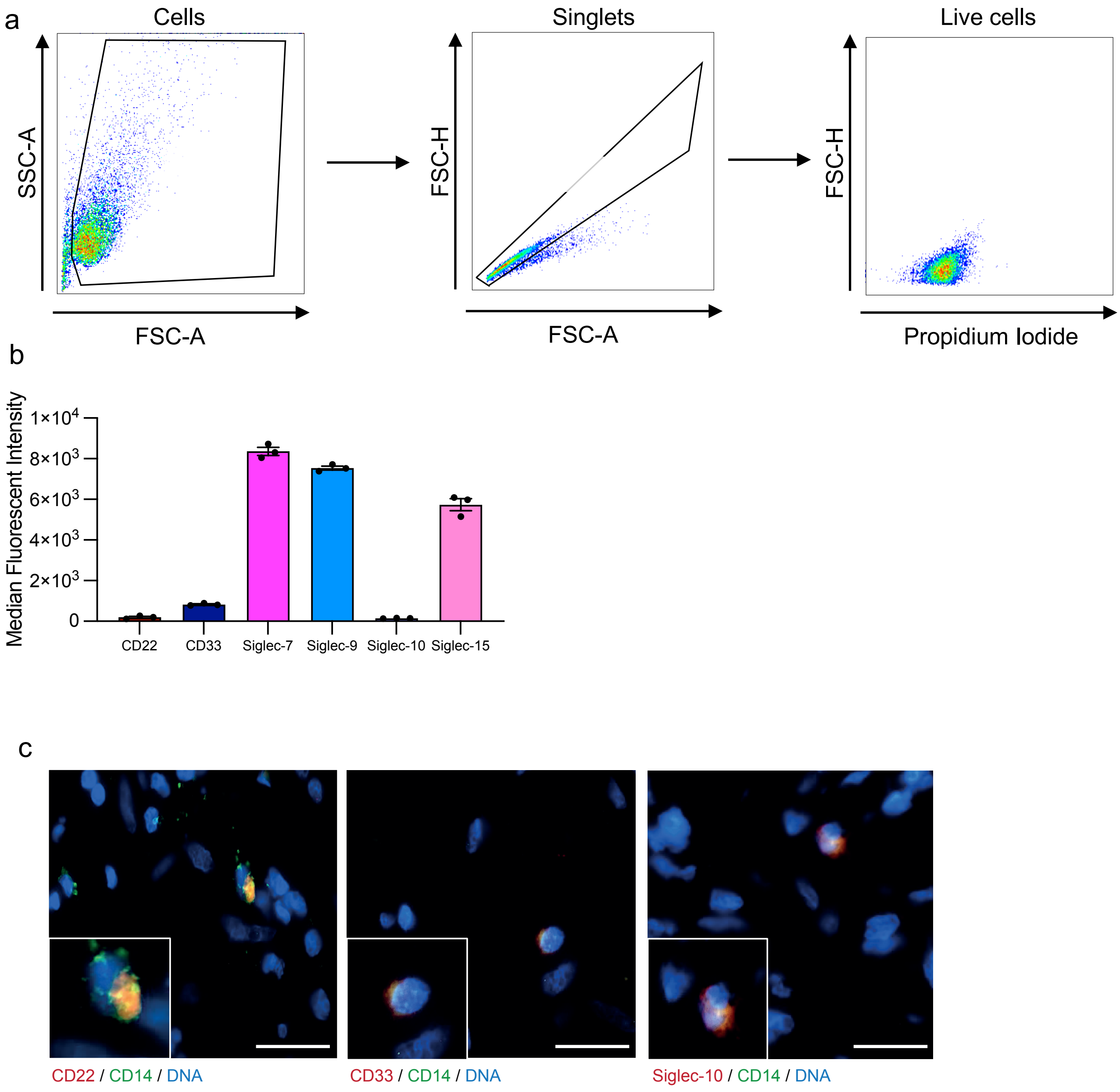

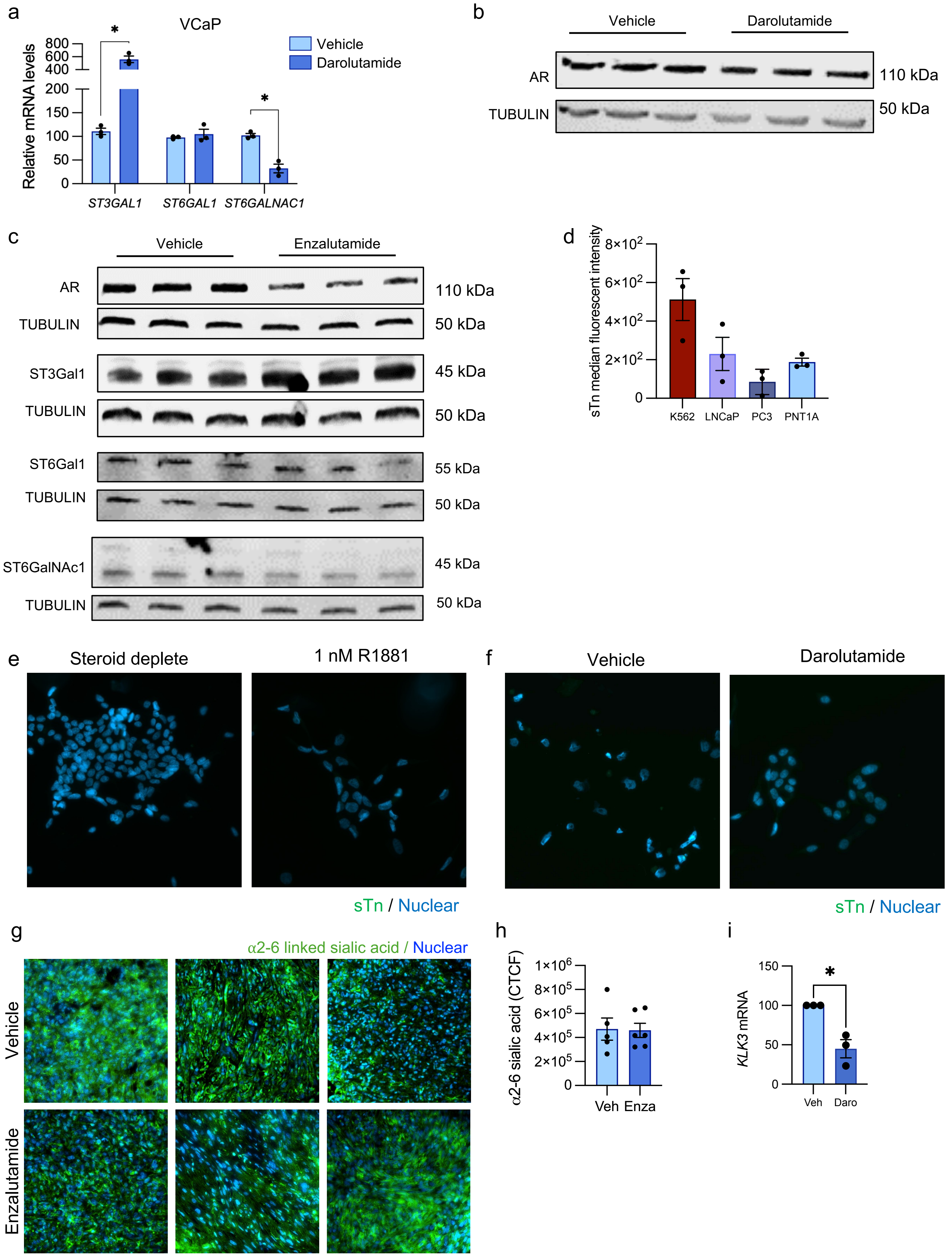

a

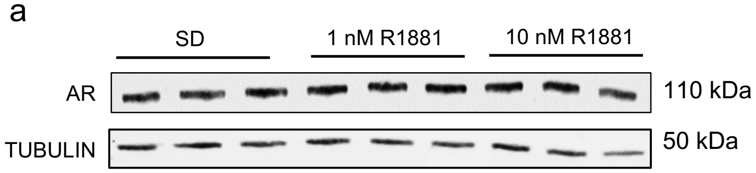

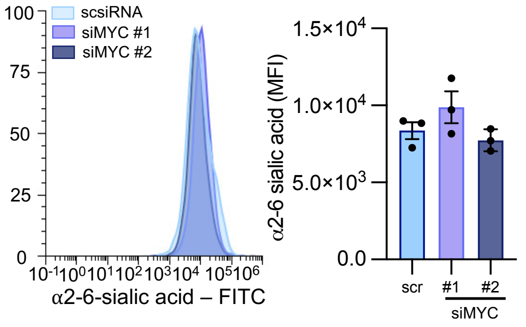

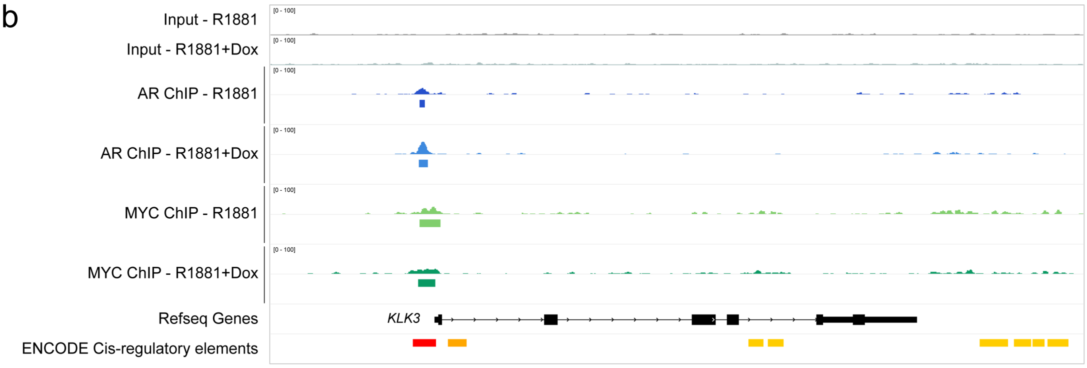

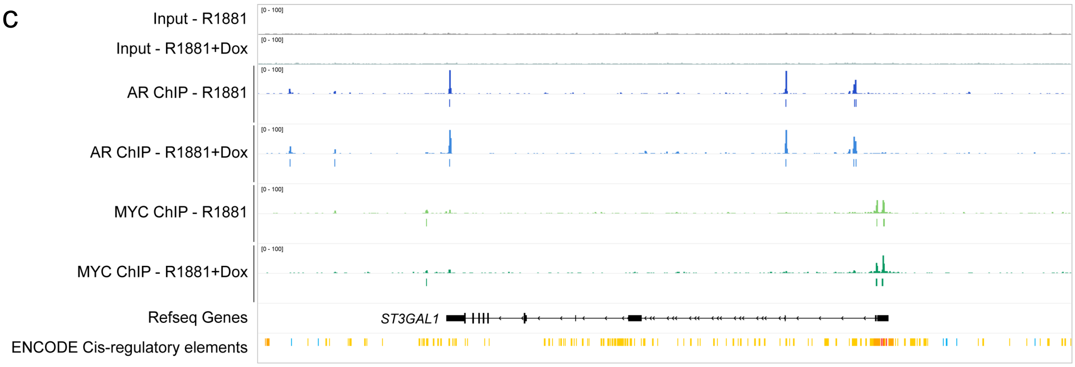

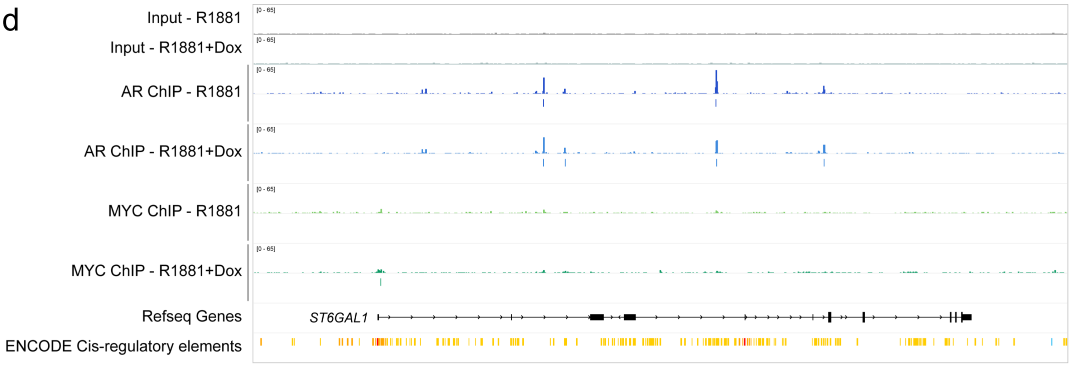

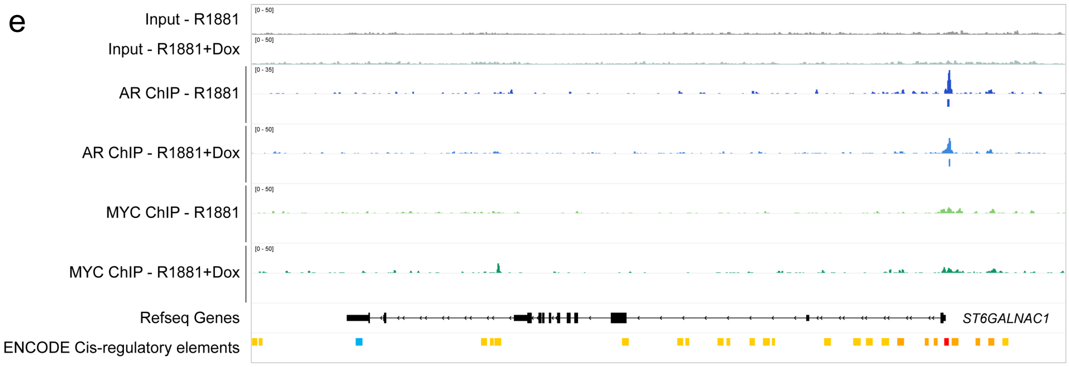

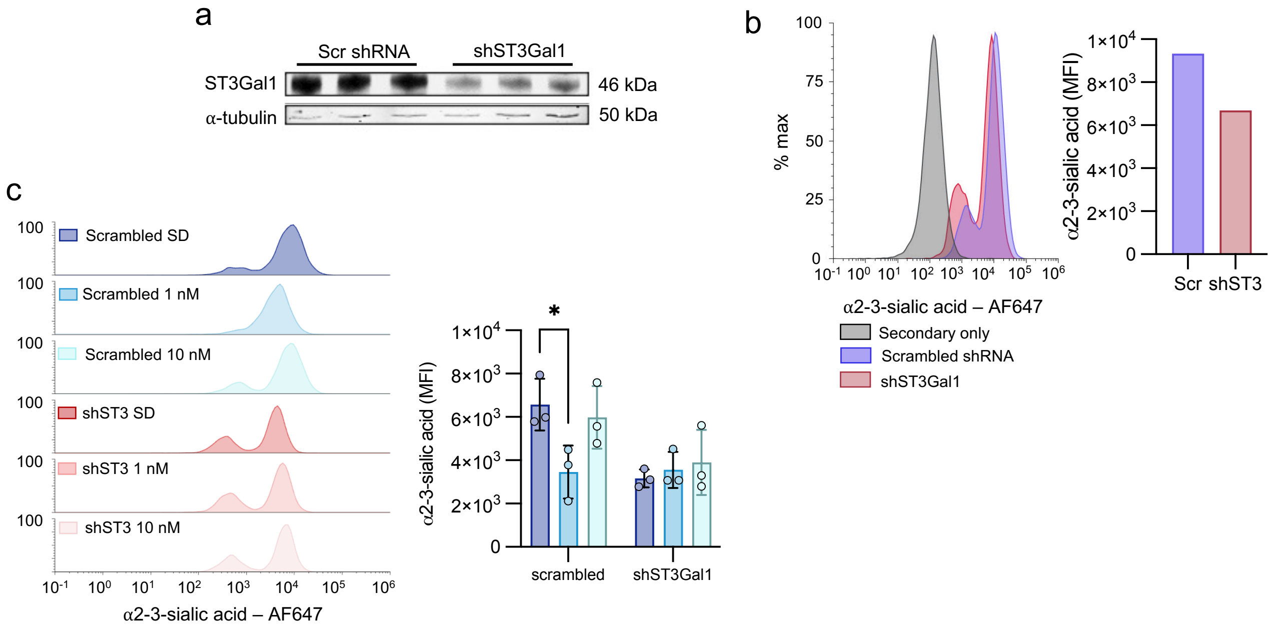
