## Supplemental Table 1 for "The Androgen Receptor and MYC synergise to modulate the synthesis of Siglec-7 ligands in prostate cancer"

Supplementary Table 1. Antibodies/Lectins

| Target | Host Species | Dilution | Manufacturer | Product No. | Application |
|---|---|---|---|---|---|
| ST3Gal1 | Rabbit | 1/500 | Invitrogen™ | PA5-21721 | Western Blot |
| Anti-hST3Gal1 | Sheep | 1/200 | R & D Systems™ | AF6905 | Western Blot |
| ST6Gal1 (Center) | Rabbit | 1/1000 | ABGENT™ | AP19891c | Western Blot |
| ST6GalNAc1 | Rabbit | 1/500 | Protein Tech® | 15363-1-AP | Western Blot |
| c-MYC | Rabbit | 1/6000 | Protein Tech® | 10828-1-AP | Western Blot |
| Androgen Receptor | Rabbit | 1/1000 | Protein Tech® | 22089-1-AP | Western Blot |
| Anti-α-tubulin | Mouse | 1/4000 | Sigma-Aldrich® | T9026 | Western Blot |
| IRDYE® 800CW Goat Anti-Rabbit IGG Secondary Antibody | Anti-Rabbit | 1/20,000 | LICORbio™ | 926-32211 | Western Blot |
| IRDYE 680RD Goat Anti-Mouse IGG Secondary Antibody | Anti-Mouse | 1/20,000 | LICORbio™ | 926-68070 | Western Blot |
| Peroxidase-conjugated AffiniPure® Goat Anti-Rabbit IgG (H+L) | Anti-Rabbit | 1/4000 | Jackson ImmunoResearch™ | 111-035-003-JIR | Western Blot |
| Peroxidase-conjugated AffiniPure® Goat Anti-Mouse IgG® (H+L) | Anti-Mouse | 1/4000 | Jackson ImmunoResearch™ | 115-035-003-JIR | Western Blot |
| Rabbit Anti-Sheep IgG (H+L) Secondary Antibody, HRP | Anti-Sheep | 1/4000 | Invitrogen™ | 31480 | Western Blot |
| AMACR | Rabbit | 1/200 | Protein Tech® | 15918-1-AP | IF |
| Siglec-2 (CD22) | Mouse | 1/100 | Protein Tech® | 66103-1-Ig | IF |
| Siglec-3 (CD33) | Rabbit | 1/25 | Life Tech® | MAI-25911 | IF |
| Siglec-7 | Rabbit | 1/50 | Protein Tech® | 13939-1-AP | IF |
| Siglec-10 | Rabbit | 1/50 | Invitrogen™ | PA5-55501 | IF |
| Siglec-15 | Rabbit | 1/10 | Abcam™ | Ab198684 | IF |
| CD14 | Mouse | 1/50 | Protein Tech® | 66868-1-Ig | IF |
| CD163 | Rabbit | 1/500 | Abcam™ | Ab213612 | IF |
| Invitrogen™ Goat anti-Rabbit IgG (H+L) Highly Cross-Adsorbed Secondary, Alexa Fluor™ Plus 488 | Anti-Rabbit | 1/1000 | Invitrogen™ | A32731 | IF |
| Goat Anti-Rabbit IgG (H+L) Alexa Fluor® 488 | Anti-Rabbit | 1/1000 | Abcam™ | Ab150077 | IF |
| Donkey Anti-Rabbit IgG (H+L) Highly Cross-Adsorbed Secondary, Alexa Flour™ 594 | Anti-Rabbit | 1/500 | Invitrogen™ | A21207 | IF |
| Donkey Anti-Mouse IgG (H+L) Alexa Fluor® 594 | Anti-Mouse | 1/1000 | Abcam™ | Ab150108 | IF |
| Invitrogen™ Donkey Anti-Mouse IgG (H+L) Highly Cross-Adsorbed Secondary Antibody Alexa Fluor™ 647 | Anti-Mouse | 1/1000 | Invitrogen™ | A31571 | IF |
| Anti-tag-72 [B72.3 + CC49] | Mouse | | Abcam™ | Ab199002 | IHC |

| | | | | | |
|---|---|---|---|---|---|
| Rabbit Anti-Mouse Immunoglobulins/ Biotin (affinity isolated) | Anti-Mouse | 1/200 | Dako | E0354 | IHC |
| Maackia Amurensis Lectin II (MAL-II), Biotinylated | - | 1/1000 | Vector Labs™ | B-1265-1 | Flow Cytometry/ IF |
| Sambucus Nigra (Elderberry Bark) Lectin (SNA, EBL), fluorescein (FITC) | - | 1/1000 | Invitrogen™ | L32479 | Flow Cytometry |
| Streptavidin AZDye™ 647 Monovalent Antibody Labelling Kit | - | 1/1000 | Abcam™ | Ab272190 | Flow Cytometry/ IF |
| Recombinant Human Siglec-7 Fc Chimera Protein | Human | 1/20 | R & D Systems™ | 1138-SL | Flow Cytometry |
| Recombinant Human Siglec-9 Chimera Protein | Human | 1/20 | R & D Systems™ | 1139-SL | Flow Cytometry |
| Recombinant Human Siglec-15 Chimera Protein | Human | 1/20 | R & D Systems™ | 9227-SL | Flow Cytometry |
| Goat Anti-Human IgG Fc Secondary Antibody, PE, eBioscience™ | Anti-Human | 1/1000 | Invitrogen™ | 12-4998-82 | Flow Cytometry |
