## Supplemental Table 2 for "The Androgen Receptor and MYC synergise to modulate the synthesis of Siglec-7 ligands in prostate cancer"

Supplementary Table 2. Human qPCR Primer Sequences

| Target | Forward (5’-3’) | Reverse (5’-3’) |
|---|---|---|
| ***ST6GAL1*** | ACCATTCGCCTGATGAACTC | CTGGGGCTTGAGGATGTAAA |
| ***ST6GAL2*** | GGAAGAAGGCTGGITCATTC | GTGTGGTTTCATGGCAGGTC |
| ***ST6GALNAC1*** | AGGCACAGACCCCAGGAAG | TGAAGCCATAAGCACTCACC |
| ***ST6GALNAC2*** | TCTCTCACCAAGTCATCGCC | CCGGATACACTTTGGAGGGG |
| ***ST6GALNAC3*** | GCCTGCATCCTGAAGAGAAAG | GCCGCCTGTCTGTGTAGGAG |
| ***ST6GALNAC4*** | CATGAAGGCTCCGGGTCG | CACAGGAGGATGTAGACGGC |
| ***ST6GALNAC5*** | TGGCGGACCACAAGGAAGAT | GGGGCACCATGCCATAAACA |
| ***ST6GALNAC6*** | ACTACTGCAGCGGCCC | TTCTCGGTGATGAAGCGGTG |
| ***ST3GAL1*** | GACACCCACACCCCTGTATTC | CTGAAAATGGTACCAACACGGC |
| ***ST3GAL2*** | GCTTTGAGCAGGATGTTGGC | CACTGGGGCGTAGGTGAATC |
| ***ST3GAL3*** | AAGGAGAGAGTGGAGGCCG | GCCAAGGGTAGGGATGTTCC |
| ***ST3GAL4*** | CCGGGAAGACAGGTACATCG | CTGATCCCGGGAGTAGTTGC |
| ***ST3GAL5*** | GGGGCCGAGATAAGAACGTC | CGCCAAACTGACTTCATCGC |
| ***ST3GAL6*** | GCTCCTGCAGCTTTCCAAAC | TCATGGCTGGCTCACCTTTC |
| ***ST8SIA1*** | AGAGCATGTGGTATGACGGG | GGGAGATTGCATCTGTGGCA |
| ***ST8SIA4*** | AGACCTGTGCAGTTGTGGGA | CACAGGAGCTAGATTACACCTTA |
| ***ST8SIA5*** | CTGTCCTCCCTTCTCCTTGTCTAC | CCACTGGTGTCCACCTCATAC |
| ***β-ACTIN*** | CATCGAGCACGGCATCGTCA | TAGCACAGCCTGGATAGCAAC |
| ***TUBULIN*** | CTTCGGCCAGATCTTCAGAC | AGAGAGTGGGTCAGCTGGAA |
| ***GAPDH*** | AACAGCGACACCCATCCTC | AGCACAGCCTGGATAGCAAC |
